# Elevated HIV viral load is associated with higher recombination rate *in vivo*

**DOI:** 10.1101/2023.05.05.539643

**Authors:** Elena V. Romero, Alison F. Feder

## Abstract

HIV’s exceptionally high recombination rate drives its intra-host diversification, enabling immune escape and multi-drug resistance within people living with HIV. While we know that HIV’s recombination rate varies by genomic position, we have little understanding of how recombination varies throughout infection or between individuals as a function of the rate of cellular coinfection. We hypothesize that denser intra-host populations may have higher rates of coinfection and therefore recombination. To test this hypothesis, we develop a new approach (Recombination Analysis via Time Series Linkage Decay, or RATS-LD) to quantify recombination using autocorrelation of linkage between mutations across time points. We validate RATS-LD on simulated data under short read sequencing conditions and then apply it to longitudinal, high-throughput intra-host viral sequencing data, stratifying populations by viral load (a proxy for density). Among sampled viral populations with the lowest viral loads (< 26,800 copies/mL), we estimate a recombination rate of 1.5×10^−5^ events/bp/generation (95% CI: 7×10^−6^−2.9×10^−5^), similar to existing estimates. However, among samples with the highest viral loads (> 82,000 copies/mL), our median estimate is approximately 6 times higher. In addition to co-varying across individuals, we also find that recombination rate and viral load are associated within single individuals across different time points. Our findings suggest that rather than acting as a constant, uniform force, recombination can vary dynamically and drastically across intra-host viral populations and within them over time. More broadly, we hypothesize that this phenomenon may affect other facultatively asexual populations where spatial co-localization varies.

## Introduction

Recombination is a key evolutionary driver, permitting organisms to purge deleterious mutations and combine beneficial ones. These functions are critical in human immunodeficiency virus (HIV) [37] where intra-host recombination promotes viral diversification, immune escape [42], multi-drug resistance [23, 33, 36] and the maintenance of genomic integrity in spite of an exceptionally high viral mutation rate [25, 41]. While average HIV recombination rates have been previously estimated on the order of 10^*−*5^/bp/generation or higher [51, 34, 6], new investigations have revealed that this average fails to capture the full variation in recombination rate in multiple settings [48, 53, 54]. Because of recombination’s critical role in HIV’s evolutionary success, it is important to investigate the factors that underlie its variation.

Two processes govern HIV’s recombination rate: cellular coinfection and reverse transcriptase template switching (Figure 1A). When multiple viruses co-infect the same cell, the virions emerging from that cell can contain two distinct but co-packaged viral genomes (as opposed to virions produced by singly infected cells which contain two identical genomes) [22, 9]. When those virions infect new cells, template switching between the two co-packaged genomes can result in a recombinant genome inheriting alleles from both parent strands [38, 20, 19]. Increased rates of coinfection should result in more virions with distinct co-packaged genomes, in which recombination can create new viral genotypes, while increased rates of template switching should result in more recombination events per genome pair.

**Figure 1.**
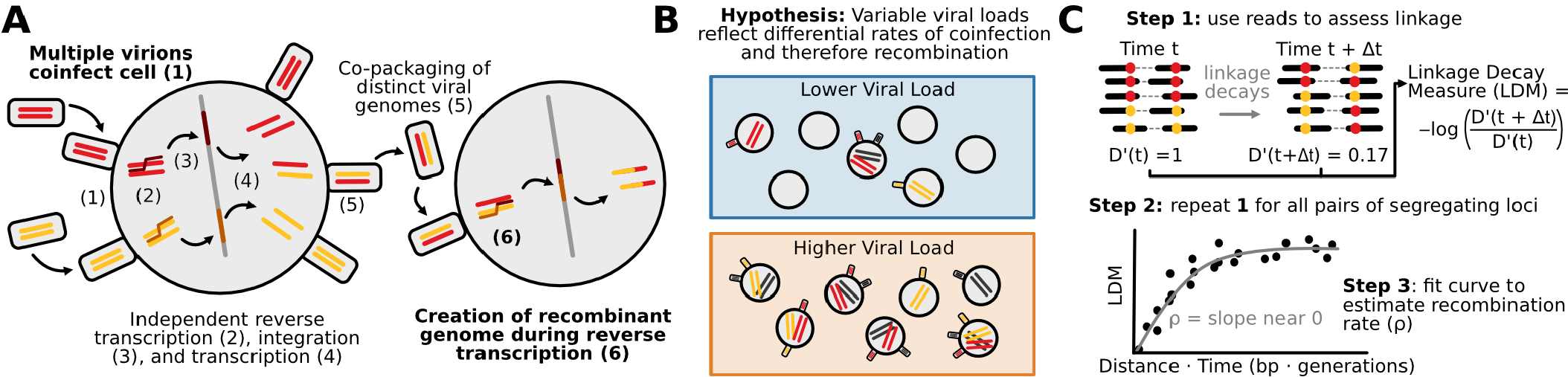
Drivers of HIV recombination rate. **(A)** Cartoon schematic of HIV replication cycle leading to recombination between distinct viral genomes. Steps that determine the rate of recombination are shown in bold. **(B)** HIV populations with higher viral loads (orange) may experience higher rates of viral coinfection and therefore higher rates of recombination than HIV populations with lower viral loads (blue). **(C)** Linkage decay patterns enable recombination rate estimation. D’ values quantify genetic linkage between pairs of SNPs at a given time point. Negative log ratios of D’ values (here termed linkage decay measures or LDMs) across two sampled time points are positively associated with the product of the time between sample points and distance between SNPs. The slope of this relationship can be used to estimate the recombination rate, *ρ*. We call this method RATS-LD.

Variation in the template switching rate has already been recognized as an important mediator of viral recombination both *in vitro* and *in vivo*. Template switching and, by extension, recombination can be suppressed by decreased sequence similarity between the two template strands [4, 3, 5, 35]. This effect is dose dependent [3, 35] and extremely low levels of sequence similarity (*<* 63%) near fully block template switching and recombination, leading to a breakdown of genomic integrity [41]. Recent sequence context aware work from human-derived HIV-1 sequences has identified particular genomic positions as hot and cold spots for template switching and recombination, which impact the ways in which viral populations diversify [32, 48, 53, 54].

Despite its potential importance, we know little about how variation in coinfection rates impacts viral recombination *in vivo*. While studies of HIV in mouse models and cell cultures have demonstrated that increased coinfection is associated with an increase in recombinant viruses [27], it is unknown if this effect persists among naturally-occurring HIV infections in humans. We expect coinfection to be mediated by the abundances and locations of both viruses and the CD4+ T cells they infect [22, 9]. Therefore, we expect viral load - a proxy for overall viral abundance - to be related to the rate at which coinfection occurs, even if non-linearly. While previous studies have found a weak relationship between viral load and coinfection rate in chronic infection, it is unclear how generalizeable this relationship is and what effect, if any, it has on viral recombination rate [21]. Since viral load and CD4+ T cell count vary across individuals and within them over time, we hypothesize that viral intra-host demography modulates coinfection rates and may therefore play an important but previously uncharacterized role in determining the viral recombination rate in human hosts (Figure 1B).

Existing statistical approaches to estimate recombination rate have made this effect challenging to detect *in vivo*. Current approaches to estimate the HIV recombination rate fall into two broad categories: identifying specific breakpoints in sequencing data [52, 31, 54, 30], or employing overall patterns of linkage disequilibrium to infer intra-host recombination rates (breakpoint-free) [6, 34, 51]. Both rely on the analysis of single genome amplification and sequencing data [45], which substantially limits sampling depth and makes quantitative comparisons across different viral load groups challenging. Note, Song et al. found no differences in recombination breakpoint counts at different viral loads, likely because the number of breakpoints visible at any specific point in time is small [54]. While bulk-sequencing data enables deeper sampling than single genome sequencing, the relatively short reads commonly used in bulk-sequencing protocols limit linkage measurement except between nearby sites. However, HIV’s high recombination rate results in significant breakdown of genetic linkage even within the relatively short length of a typical bulk sequencing paired-end read. Therefore, adapting breakpoint-free methods to work with bulk sequencing data has the potential to more sensitively quantify the effects of recombination, even when stratifying data sets by viral load.

Here, we test for viral load-associated differences in recombination rate through developing a new approach called Recombination Analysis via Time Series Linkage Decay (RATS-LD) which quantifies recombination rates from longitudinal bulk sequencing data. We validate that this method can successfully recover recombination rates in simulated short read paired-end data. To test for positive associations between HIV viral loads and recombination rate, we apply RATS-LD to longitudinal intra-host HIV bulk sequencing data from individuals with varying viral loads sequenced with careful controls for PCR recombination [56, 55]. We find that while HIV populations with viral loads in the lowest third of data set (*<* 26, 800 copies/mL) have recombination rates in line with previous estimates (1.5*×*10^*−*5^/bp/generation), populations with viral loads in the upper third (*>* 82, 000 copies/mL) have a median recombination rate that is nearly six-fold higher. Furthermore, within single individuals we observe patterns of viral load and effective recombination rate increasing concurrently. These findings demonstrate that recombination rate is not a static parameter, and intra-host conditions can mediate the strength of its effect both across and within individuals over time. Since recombination mechanisms generally require that two genomes physically find each other in space, such contact-network mediated effects are also likely to occur in populations beyond HIV.

## Results

### Measuring intra-host recombination rates using short read data

We developed an approach to leverage short read sequencing data to quantify recombination rates in longitudinally-sampled intra-host viral populations. Our approach exploits the decay in genetic linkage following the emergence of new mutations in the population. New mutations will initially occur on a single genetic background and will be measurably linked to any other single nucleotide polymorphisms (SNPs) on that background. We measure this linkage using the *D*′ statistic [28] which is given as follows:

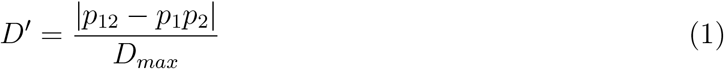

where *p*_1_ and *p*_2_ are the frequencies of the majority nucleotide at the corresponding locus, *p*_12_ is the frequency of the haplotype made up by the majority nucleotides at the two loci, and *D*_*max*_ serves as a normalizing factor (See Materials & Methods for details on *D*_*max*_ calculation). This statistic captures the difference between the observed frequency of the majority haplotype and its expected frequency, *p*_1_*p*_2_, if the loci were unlinked. When SNPs are maximally linked, D’ = 1, and when they are at linkage equilibrium, D’ = 0. Over time, recombination will disassociate linked groups of mutations, leading to a relaxation back towards linkage equilibrium. This disassociation will be faster if the SNPs are separated by greater physical distances (*d*, measured in nucleotides), as the increased distance provides more opportunities for recombination events between the SNPs. Specifically, for a pair of segregating SNPs, the linkage decay after a period of time Δ*t* with a recombination rate *ρ* is given as follows [18]:

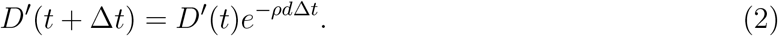

Thus, by examining the genetic association between pairs of SNPs longitudinally, we can quantify the rates of this statistical decay at different distances to estimate *ρ*:

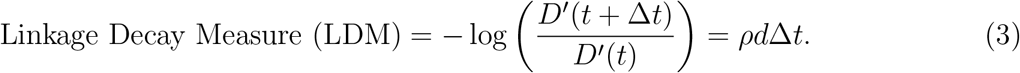

We compute the negative log ratio on the left hand side of equation 3 (which we call the linkage decay measure or LDM) over many pairs of SNPs between multiple time points. We can then use these LDMs to derive an empirical relationship between the linkage decay and the time and distance separating each LDM’s corresponding pair of SNPs (Figure 1C). The initial slope of this empirical relationship is the estimated recombination rate 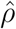. This approach, which we call Recombination Analysis via Time Series Linkage Decay (or RATS-LD), is a generalization of existing methods for recombination rate estimation [34] which permits application in short read sequencing data.

### Validating RATS-LD on simulated data under neutrality

We first simulated neutrally evolving intra-host viral populations with known recombination rates and tested that RATS-LD could recover these known rates. To create input for RATS-LD, we sampled these simulations longitudinally and mimicked the short-read, paired-end sampling common in viral deep sequencing experiments. We centered our simulated recombination rates around existing HIV intra-host estimated recombination rates (*ρ* = 2*×*10^*−*6^ - 10^*−*3^ [6, 34, 51], see Materials & Methods for full simulation and sampling details). In these simulations, we observed that the relationship between the linkage decay measure and the time-scaled physical distance (*d*Δ*t*) is not linear, but asymptotically approaches a given LDM at large *d*Δ*t* values. This relationship emerges because, at long time-scaled distances, the genetic association between two SNPs becomes dominated by other evolutionary forces and our resolution to measure linkage decay is overtaken by noise. The rate at which this asymptote is approached depends strongly on the recombination rate (Figure 2A, Supplemental Figure S1).

**Figure 2.**
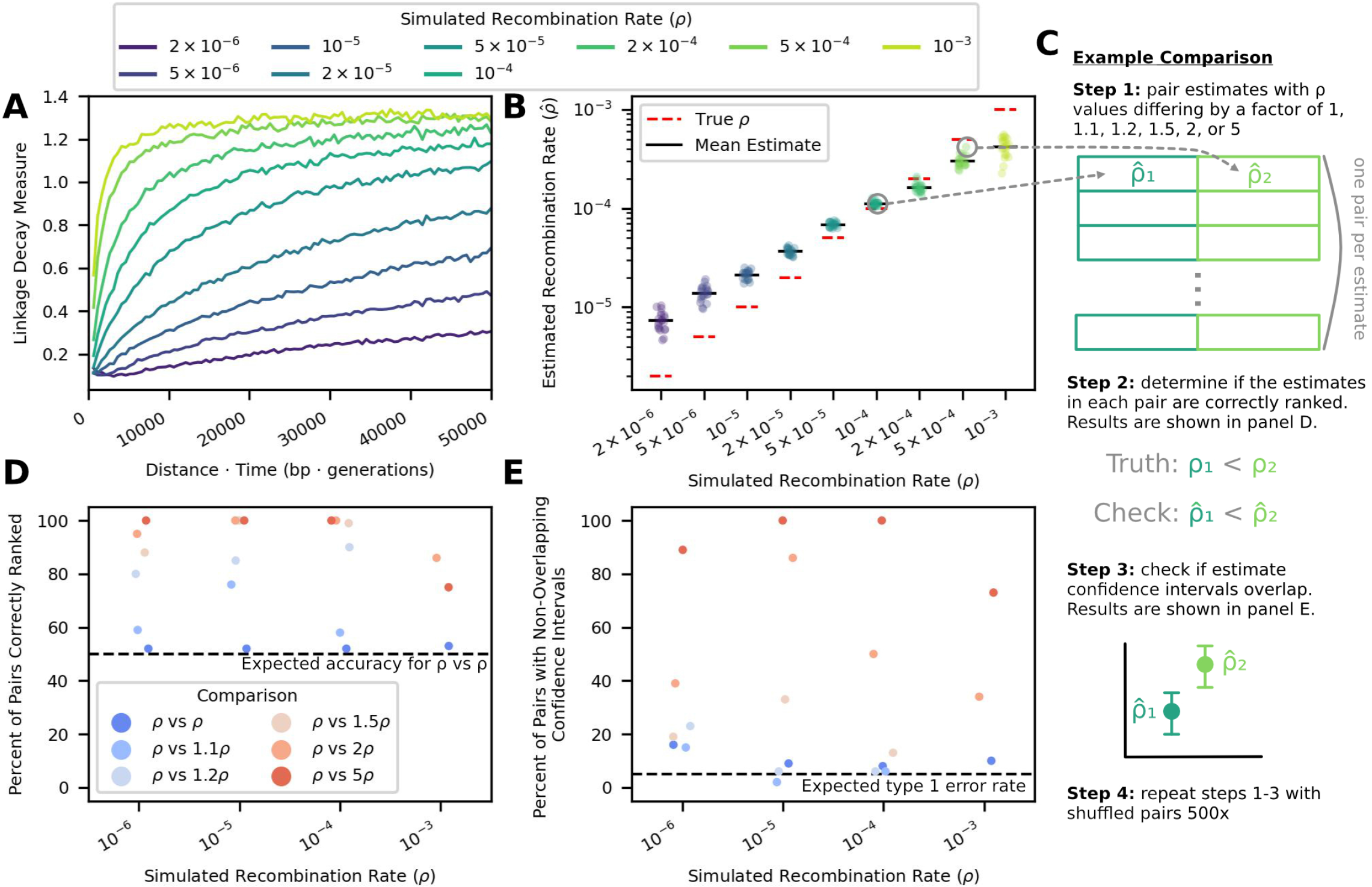
RATS-LD recaptures true recombination rates in simulated data and accurately distinguishes between different rates. **(A)** Linkage decay curves calculated from simulated data exhibit saturation patterns with respect to *d*Δ*t* that depend on the underlying recombination rate, *ρ* (moving average displayed, window size = 500 bp generations). **(B)** The slopes of these curves at *d*Δ*t* = 0 serve as RATS-LD’s estimates and recapture the known *ρ* value in simulated data. Each point represents the median of a bootstrapped distribution of 1000 estimates. **(C)** We performed a discrimination analysis to confirm that RATS-LD can distinguish different recombination rates. We simulated data with set recombination rates and estimated 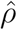 using RATS-LD. We then randomly paired simulations with recombination rates that differed by a factor of 1, 1.1, 1.2, 1.5, 2, or 5 to assess how often RATS-LD correctly distinguished underlying recombination rates via simple rank order **(D)** and non-overlapping bootstrapped 95% confidence intervals **(E)**.

To account for this asymptotic effect, we modified our approach for estimating *ρ* by fitting a function of the form

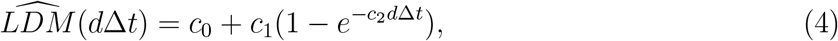

which can capture this behavior. The derivative of 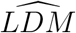 with respect to *d*Δ*t* at 0 recaptures the true recombination rate in simulated data from neutrally-evolving populations (Figure 2B). We noted that estimation at high or low values of *ρ* was biased downward or upward, respectively.

This minor bias emerged not because the linkage decay measure and the time-scaled distance scale unexpectedly, but due to challenges fitting linkage decay curves with limited paired-end data. To demonstrate that the slope of the linkage decay measure at 0 does indeed recapture the true recombination rate, we fit linear relationships between *d*Δ*t* and the linkage decay measure for subsets of the data near *d*Δ*t* = 0 (i.e., pre-curve saturation). We found that these linear fits accurately recaptured the true recombination rate when only pre-saturation data was included in the fit (Figures S2 & S3). However, the linkage decay curves saturate at different *d*Δ*t* for different values of *ρ*, rendering these linear fits unsuitable when the true recombination rate is unknown (Figures 2A & S3). In contrast, the curve-fitting approach can be applied without a known *d*Δ*t* saturation point. For low recombination rates, the linkage decay curves do not saturate in the time-scaled distances available from the relatively short simulated reads (all pairs of SNPs less than 700 bp apart) (Figure 2A). As a result, RATS-LD underestimates the asymptote of Equation 4, which results in a slight overestimation of *ρ* (Figure 2B). Among very high recombination rates, little data is available before curve saturation, resulting in challenges fitting Equation 4’s behavior near *d*Δ*t* = 0 and underestimation of *ρ*. For full details on how these fits were performed, see Materials & Methods. See the discussion for a description of best practices for assessing whether or not a RATS-LD fit should be considered reliable.

Despite the challenges of quantifying the recombination rate at more extreme *ρ* values, we found that RATS-LD could readily discriminate between differences in recombination rates in simulated data, particularly in the range of previously estimated HIV recombination rates (10^*−*5^*−*10^*−*4^). We assessed this in two ways: first, we compared RATS-LD estimates (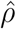 values) across pairs of simulations and checked if the simulation with the higher true *ρ* also had the higher estimated 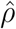 (Figures 2C & 2D). In the range of 10^*−*5^ to 10^*−*4^, we found that pairs of simulations with a 1.5-fold or greater difference in *ρ* were nearly always ordered correctly using RATS-LD. Outside of this range (at values of 10^*−*6^ and 10^*−*3^), recombination rate differences must be larger for RATS-LD to successfully order them due to lower method resolution in these regimes. Second, for each pair of simulations, we checked whether their bootstrapped 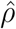 confidence intervals overlapped (Figure 2E). We found that we had a low probability of type I errors (i.e., assessing differences in recombination rates when no such differences existed). At recombination rates of 10^*−*3^ and 10^*−*6^, RATS-LD could often discriminate simulations with true *ρ* values separated by a factor of 2 or 5. In the range of 10^*−*5^ to 10^*−*4^, RATS-LD could nearly always discriminate simulations with true *ρ* values a factor of 5 apart and could often discriminate *ρ* values a factor of 2 apart, but was not well-powered to detect more subtle differences in recombination rate.

### Validating RATS-LD under settings that better recapitulate intra-host HIV

While RATS-LD can discriminate between recombination rates in neutrally-simulated populations, intra-host HIV is under strong selective pressures. We therefore extended our simulations to mimic these pressures, adding both positive and negative selection (*s* ranges from -0.1 to 0.05, see Materials & Methods for full distribution of fitness effects). We chose these values based on estimates from the literature [57], and confirmed that they match *in vivo* intra-host divergence and diversity over the first 8 years of infection in a longitudinally-sampled cohort ([56], Figure S4). We found that selection simulations produced noisier linkage decay curves than neutral simulations (Figure S5), but RATS-LD was still able to recapture underlying recombination rates (Figure 3A). Similar to the neutrally-simulated data, RATS-LD successfully estimated recombination rates in the 10^*−*5^*−*10^*−*4^ range but lost resolution at more extreme values.

**Figure 3.**
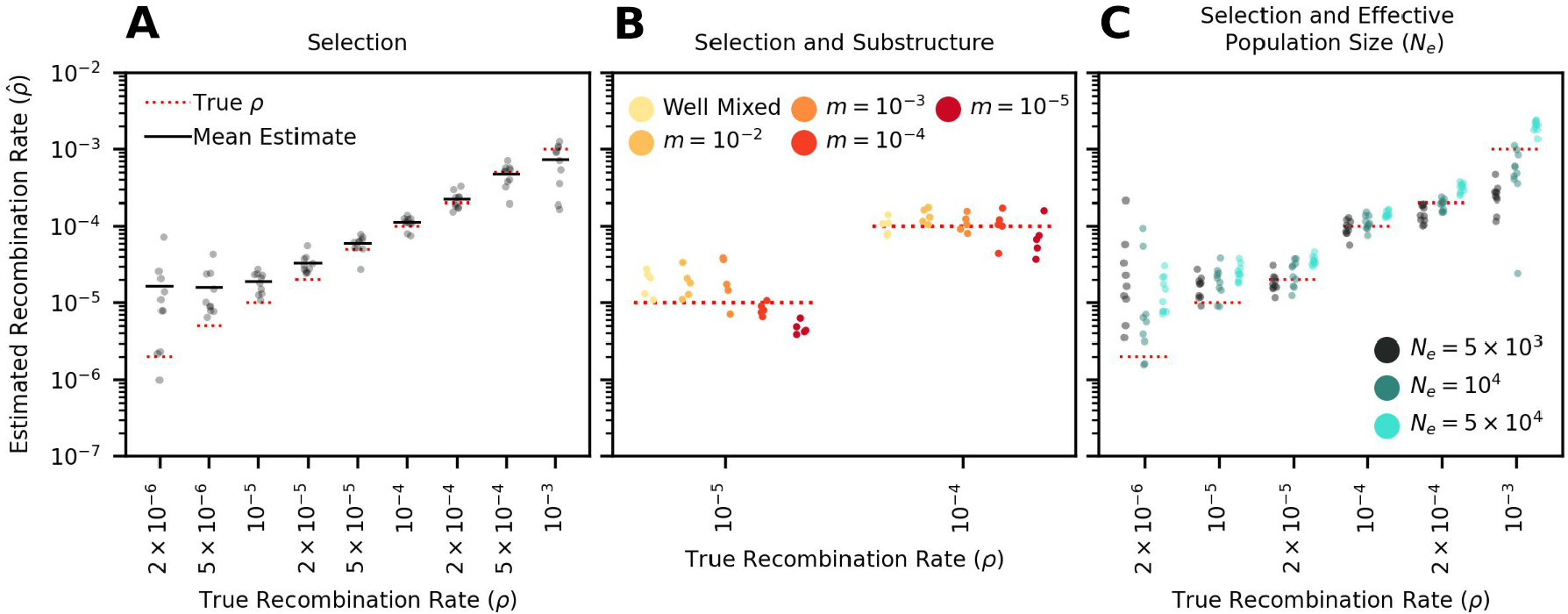
RATS-LD can estimate recombination rates in conditions that mimic HIV intra-host evolution. **(A)** Recombination rate estimates from simulations performed with HIV-like selective conditions recapture known *ρ* values. **(B)** Recombination rate estimates from simulated subdivided populations interconnected by bidirectional migration. Each generation, a proportion *m* of each subpopulation emigrates from the other subpopulation (*m* = 10^*−*2^, 10^*−*3^, 10^*−*4^, 10^*−*5^). **(C)** Recombination rate estimates in data sets simulated under three different effective population sizes (*N*_*e*_ = 5*×*10^3^, 10^4^, 5 10^4^). In all three panels, each point represents the median estimate over 1000 bootstraps. For full details on all of the specific simulation scenarios see Materials & Methods.

Another complication of intra-host HIV evolution is population structure between different viral subpopulations in the body, which can maintain longer range linkage and potentially affect estimation. To examine the impact of population structure, we simulated two sub-populations of HIV interconnected by migration at rate *m*. We chose a range of *m* values to broadly encompass extremely low migration rates considered in the literature [29] up to well-mixed populations.

We found that RATS-LD estimation was minimally impacted by migration rates greater than 10^*−*4^, although extremely low migration rates (*m* = 10^*−*5^) did depress estimates slightly (Figure 3B). To understand if this underestimation was likely to impact application of RATS-LD to *in vivo* data with an unknown amount of population substructure, we compared the average linkage in the simulated structured data to the *in vivo* data from Zanini et al. [56] used above as a reference for genetic divergence and diversity. Notably, this data set contained two superinfected individuals who displayed long range linkage suggestive of population structure. Even in these individuals, D’ decayed faster over distance than the simulated *m* = 10^*−*5^ populations (Figure S6). This suggests that these participants did not possess migration rates low enough to substantially alter RATS-LD’s performance.

Finally, we considered whether variation in effective population size (*N*_*e*_) would change linkage patterns strongly enough to alter RATS-LD’s accuracy. We therefore ran selection simulations for two additional values of *N*_*e*_ (a factor of 2 lower and 5 higher) consistent with variation in estimated short-term *N*_*e*_ of intra-host HIV in the literature [11, 2]. We found that these changes to the effective population size altered the linkage patterns enough to affect estimated *ρ* values (Figure 3C). However, within the range of recombination rates in which RATS-LD works well, these effects were relatively small. For example, the average estimated recombination rates for *N*_*e*_ values a factor of 10 apart were different by less than a factor of 2 (at *ρ* = 2*×*10^*−*5^/bp/generation, a rate near the prevailing estimate [34]).

### Applying RATS-LD to *in vivo* HIV data

Thus established, we sought to use RATS-LD to quantify how recombination rate varied within and across intra-host HIV populations with different viral loads. We analyzed viral sequences from a cohort of 10 people living with HIV, who were sampled longitudinally over 6-12 years and were not on antiretroviral therapy (ART) [56], using a workflow illustrated in Figure S7. We chose this data set because the authors recorded contemporaneous viral load measurements and sequenced these data via a workflow that minimized PCR recombination (see Discussion) [55]. Therefore, we expect the detected signals of recombination in these data to be driven by *in vivo* processes, rather than amplification artifacts. This data set also had a sampling depth (median *≈*800) far deeper than was necessary for accurate RATS-LD estimation (Figure S8A). RATS-LD estimated the average recombination rate across this data set to be 3.6*×*10^*−*5^ events/bp/generation (95% CI 2.8*×*10^*−*5^*−* 4.7*×*10^*−*5^) which is slightly higher than previous estimates [6, 34, 51], but consistent under RATS-LD’s slight upward bias in this range.

Within this data set, viral loads varied within and between individuals (Figure S9), as has been observed in other cohorts [43, 50]. For each pair of time points within an individual, we computed the mean viral load (Figure S10A). Note, time points did not need to be consecutive to be compared, but we excluded any time point pairs where an intervening viral load measurement did not fall between the measurements at the two compared time points (see Materials & Methods). Next, we associated each linkage decay measure to the mean viral load between the two time points (*t* and *t* + Δ*t*). The distribution of LDMs and their associated average viral load measures is shown in Figure S10B. Then, we split the data based on the viral load tertiles which divided the data into three groups with approximately equal numbers of LDMs. This provided a sufficient number of loci for estimation (Table S1 & Figure S11B-C). In Figure 4A, we show the viral load time courses for each participant with shading corresponding to the viral load tertiles.

**Figure 4.**
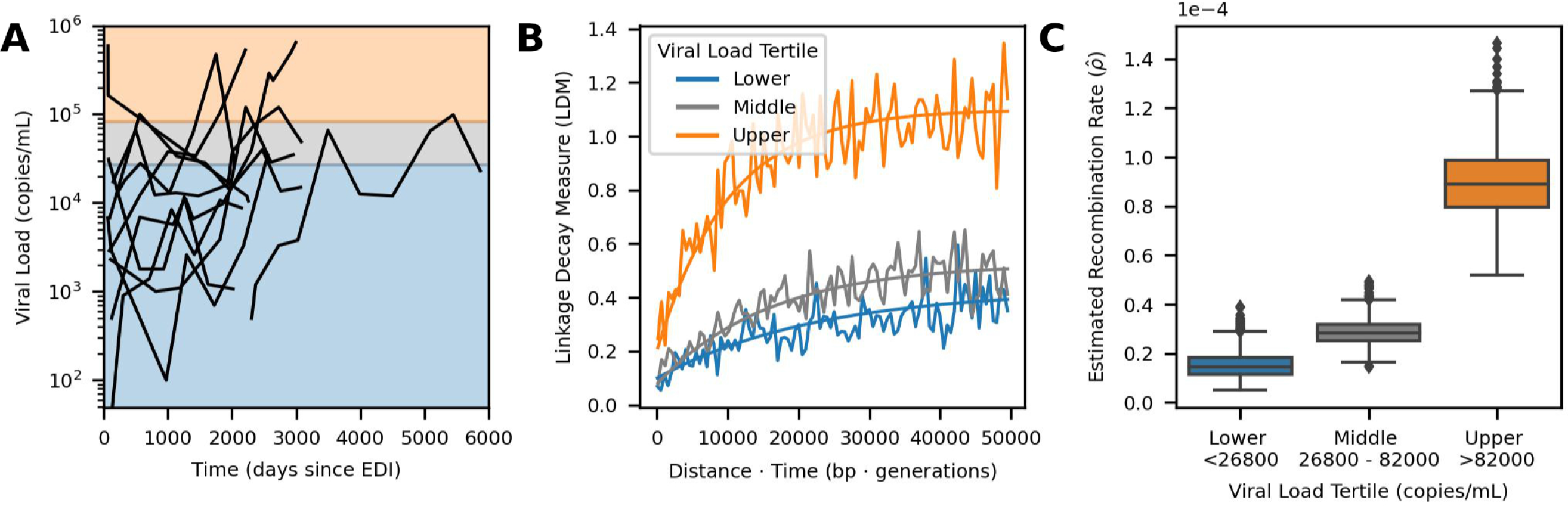
Elevated recombination rates are associated with high viral load *in vivo*. **(A)**Viral load time courses for each participant are plotted. The background shading indicates the viral load tertiles: *<* 26, 800 (blue), 26, 800*−* 82, 000 (gray), and *>* 82, 000 (orange) which divide the linkage decay measures (LDMs) into three approximately equal sized groups (*≈* 20k LDMs per group). While viral loads at single time points are shown here, pairs of time points were used to form the groups for estimation and were sorted based on their average viral load (see Figures S9 & S10). **(B)** Curve fits used by RATS-LD for estimation. Curve fits representing the median estimates of the bootstrap distribution (1000 bootstraps) for each group are overlaid on the binned average of the LDMs from the *in vivo* data (moving average used for display purposes only, window size = 500 bp · generations). **(C)** Recombination rate estimates 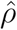with 1000 bootstraps are shown separately for the three tertiles. Each box represents the quartiles of the 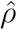distribution and the median is indicated by the central line.

We used RATS-LD to compute 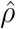 separately for each of the viral load groups (Figure 4B-C). We found that 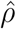 among time point pairs below the first tertile (*<* 26, 800 copies/mL) was 1.5 *×* 10^*−*5^ recombination events/bp/generation (bootstrapped confidence intervals 7 *×* 10^*−*6^ *−* 2.9 *×* 10^*−*5^), a rate nearly identical to existing recombination rate estimates for HIV [34]. However, among pairs of time points with the highest average viral loads (*>*82,000 copies/mL), the estimated median recombination rate was nearly 6 times higher: 8.9*×*10^*−*5^ recombination events/bp/generation (bootstrapped confidence intervals 6.5*×*10^*−*5^*−*1.2*×* 10^*−*4^). The middle tertile fell between the other two, although substantially closer to the lower tertile (bootstrapped confidence intervals *×*10^*−*5^*−*4*×*10^*−*5^).

We performed several analyses to ensure the robustness of this result. First, we demonstrated that the observed differences were not solely driven by the data of any individual participant by conducting a leave one out analysis (Supplemental Figure S12). Second, we performed a variation of this analysis where we left out data from two superinfected individuals (Participants 4 and 7). These individuals exhibited long range linkage patterns that are potentially consistent with population substructure maintained by limited migration, although the degree of linkage does not suggest that we are outside of the range of population structure where RATS-LD can estimate accurately (Supplemental Figure S6, Figure 3B). Regardless, the results from this analysis were also consistent with the findings displayed in Figure 4 and showed a significant difference between the lower and upper viral load tertile estimates (Supplemental Figure S12). Third, we attempted to verify this result with a biological replicate from a different data set [7]. Caskey et al. sequenced HIV from longitudinally-sampled viral populations rebounding after an infusion of broadly neutralizing antibodies. As a result of this context, populations possessed relatively low viral loads (12766 *±*9859 copies/mL, mean *±*SD). The amount of data available and low sampling depth permitted only a single, noisier estimate as opposed to the viral load-segregated analysis of Zanini et al. [56]. However, the estimate from Caskey et al. [7] was consistent with the lowest tertile data from Zanini et al. [56], in line with their comparable viral loads (Figure S8B).

Fourth, to further quantify the relationship between viral load and estimated recombination rate, we also conducted an analysis in which we grouped LDMs depending on their average viral load meeting a given threshold (Figures S13 & S14). This permitted us to further determine if viral load has a dose-dependent effect on recombination rate. The estimated recombination rate became more extreme at higher viral load thresholds, and time points with a mean viral load above 200,000 copies/mL showed an estimated 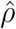 nearly an order of magnitude higher than prevailing estimates (1.1*×*10^*−*4^, confidence interval 7.5*×*10^*−*5^*−* 1.6*×*10^*−*4^). Note, however, small sample sizes led to substantial variation in curve fitting and correspondingly large bootstrapped confidence intervals at more extreme thresholds. Taken together, these results suggest that HIV populations at high viral loads may be recombining significantly more often than previous estimates.

The analyses above suggest that higher viral loads are correlated with faster recombination across individuals, but viral loads can also vary significantly within individuals over time. We therefore set out to examine if RATS-LD could detect differences in recombination rates within single individuals. Only one individual (Participant 1) in our data set had enough data above the threshold at which we saw substantial differences in 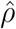 (*>*50k copies/mL, Figure S13) to perform recombination rate estimation according to our power analysis. Participant 1 possessed both extensive genetic variation (*≈* 12,000 LDMs) and a broad range of viral load measurements (*<* 10^3^ to *>* 10^6^ copies/mL). We divided the LDMs for Participant 1 into two equally sized groups based on their viral loads and found a significant difference in recombination rates between time points with viral loads above and below the median of 165,500 copies/mL (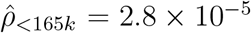, confidence interval = 1.7 *×* 10^*−*5^ - 4.5 *×* 10^*−*5^ and 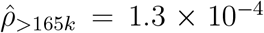, confidence interval = 7.4 *×* 10^*−*5^ - 2 *×* 10^*−*4^) as shown in Figure 5. Although we did not expect to be well-powered to detect recombination rate differences in any other participants, we repeated this analysis with Participants 3, 4, and 7, who had the next largest numbers of LDMs available for estimation (Tables S2 & S1). Consistent with the results from Participant 1, time points with viral loads below the median had lower estimated recombination rates than those with viral loads above the median (Figure S15), although the bootstrapped confidence intervals between the high and low viral load groups overlapped in all three participants. These results are not unexpected as Participants 3, 4, and 7 had much narrower ranges of viral loads than Participant 1 and RATS-LD is not well powered to detect small differences in recombination rate (Figure 2) especially at smaller sample sizes (Supplemental Figure S11). Nevertheless, the analyses within individuals collectively suggest that as a population’s viral load varies over time, evolutionarily important parameters such as the recombination rate can vary concurrently.

**Figure 5.**
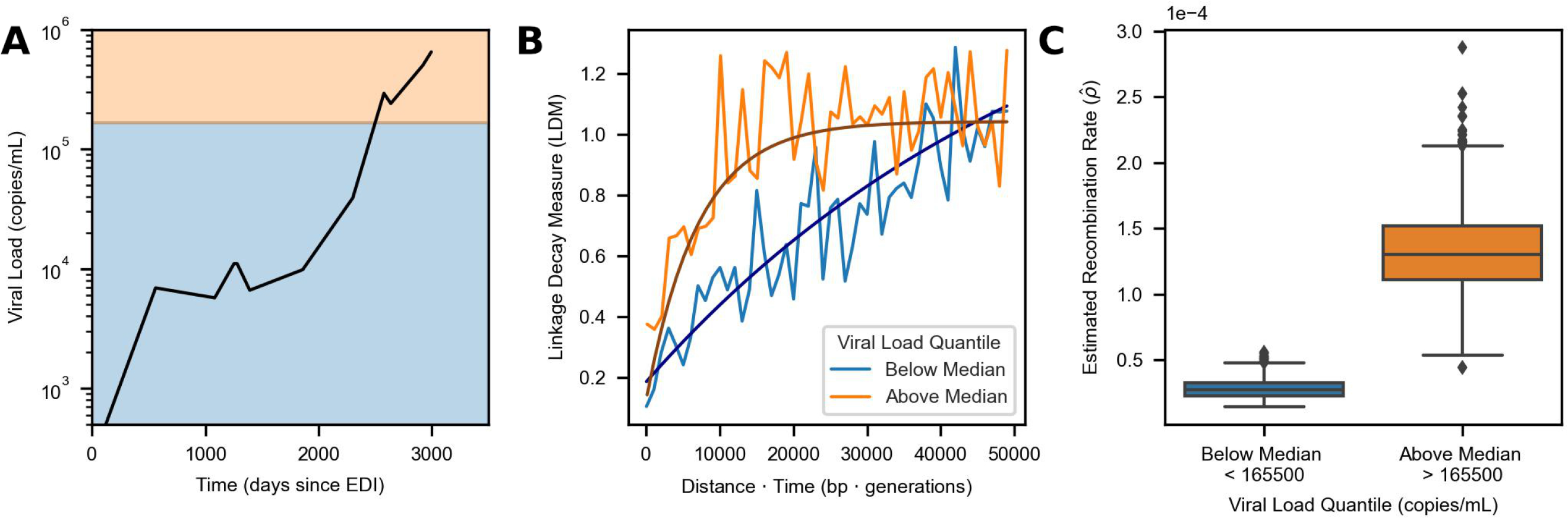
High viral load is also associated with elevated recombination rate in Participant 1. **(A)** Viral load trajectory over the number of days since estimated date of infection (EDI) for Participant 1. The background shading indicates the two groups (above median in orange and below median in blue) that pairs of time points were divided into based on their average viral load. Each group contains approximately 6, 000 LDMs. **(B)** Curve fits representing the median estimates from each group’s corresponding bootstrap distribution (1000 bootsraps) are overlaid on the binned average of the LDMs in the *in vivo* data for Participant 1 (moving average used for display purposes only, window size = 1000 bp · generations). **(C)** Recombination rate estimates for times with average viral load measurements above or below the median using only data from Participant 1. Each box indicates the interquartile range of the bootstrapped distribution of 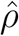 for the given viral load group (1000 Bootstraps). The median estimate is indicated via the central line. Identical analyses for Participants 3, 4, and 7 are shown in Figure S15.

## Discussion

HIV evolution within a host is a complex process driven by shifting selection pressures and variation in mutational availability as viral loads expand and contract over orders of magnitude. Here, we show that the recombination rate of HIV *in vivo* can also change dynamically over the course of infection in concert with changes in the viral load and effective population density. We introduce an approach, RATS-LD, which estimates significantly elevated recombination rates in viral populations with the highest viral loads. In addition to quantifying differences across viral populations, we also detect a nearly order of magnitude difference in recombination rates within a single viral population as its viral load increases over time. Although HIV recombination has long been appreciated to be a common event, these results suggest that in certain intra-host contexts, rates of recombination can be even more extreme than previously recognized [6, 34, 51].

Our simulations with varying effective population sizes (*N*_*e*_) suggest that these higher recombination rates among higher viral load populations are not simply caused by positive correlations between *N*_*e*_ and viral load. We found that changing HIV *N*_*e*_ within a plausible range of values resulted in much smaller differences in estimated recombination rates than the ones we observed between high and low viral load groups *in vivo* (Figure 3B). That being said, an increase in *N*_*e*_ in populations at high viral loads may be one contributing factor to the increased estimate of *ρ*. We find that recombination rate increases most drastically in populations with the highest viral loads, consistent with a study that found only modest variation in *in vivo* CD4+ T-cell coinfection rates among viral populations with viral loads less than 10^5^ copies/mL, but elevated coinfection rates in a population with viral load above 10^6^ copies/mL [21]. Our study not only suggests that these results hold in an independent cohort, but further links variable rates of coinfection to quantitatively different viral recombination rates *in vivo*. While this effect has previously been demonstrated in cell culture where viral density can be experimentally manipulated [27], this represents the first evidence that intra-host viral density varies extensively enough to cause substantial changes to the recombination rate *in vivo* in human hosts.

This previously unappreciated recombination rate variation likely impacts both intra-host evolutionary dynamics and also interpretations of intra-host viral data. While most of the sequencing data we investigated were not from acute infection, the initial HIV expansion brings the viral load its most extreme values [49]. If elevated recombination rates broadly accompany high viral loads, recombination may drive initial population diversification to a previously unrecognized extent, particularly in populations with multiple founders. The importance of this effect may be counter-acted by relatively low population diversity in the early stages of infection [56]. This interaction merits future investigation. Recent work has also suggested that elevated recombination rates could mask signatures of spatial compartmentalization in intra-host HIV populations [47]. In contrast, low viral loads in individuals on incompletely suppressive ART may lead to correspondingly low recombination rates, mitigating recombination of distinct drug resistance mutations on different genetic backgrounds as a path to multi-drug resistance (speculated in [13]). Lower rates of recombination in lower viral load populations may also make phylogenetic approaches that rely on treelike evolutionary divergence more tractable to apply without requiring ancestral recombination graphs [8]. While we did not have access to samples with extremely low viral loads near the limit of detection, future investigations could determine if previously reported average recombination rates (*≈*1.4*×*10^*−*5^) can explain low viral load patterns of linkage decay.

Viral load is the most accessible proxy for measuring population density, but it does not capture the complex dynamics of HIV metapopulations which can span distinct anatomical locations and may operate differently across individuals [29]. Since most T-cells in a person living with HIV are uninfected, local population density of infected cells is likely critically important for determining *ρ*. Future investigation into this hypothesis would be best served by localized measurements of coinfection and gene exchange in lymphatic tissues at different viral densities. These samples are invasive to collect, especially longitudinally, so approaches that consider aggregate measurements at the population level are valuable proofs of principle. The fact that positive associations with recombination rate can be observed even for high level measurements like viral load underscores density’s importance as a mediator of *in vivo* recombination.

Like other *in vivo* methods, RATS-LD relies on naturally segregating sites as genetic markers to track the breakdown of linkage. While the rate of this breakdown is not affected by population diversity, our ability to accurately quantify its average does depend on the number of segregating sites we can identify. Therefore, having more or fewer mutations stands only to impact the confidence of RATS-LD rather than bias the estimated recombination rates themselves. Less diverse populations require more pooling to have the requisite number of pairs of data points. This limits our ability to powerfully estimate recombination rates in smaller, less diverse populations.

Importantly, we cannot detect recombination events between identical sequences. If genetically identical viral bursts are largely confined to a given spatial location, rates of coinfection and recombination may be high but undetectable to this method. Similar effects could be driven by cell-to-cell transmission [26]. While important for understanding the viral life history and phenomena like complementation [16] and replication fidelity [25, 41], these undetectable recombination events are unlikely to contribute meaningfully to the population’s evolution because they do not generate new viral haplotypes. As such, our estimated 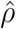 values reflect the effective rates of recombination and mixing discounting population structure.

Although we analyzed data with careful controls for PCR recombination, we cannot completely rule it out as a complicating factor [45]. In order to minimize PCR recombination, Zanini et al. only used one round of PCR (rather than a more standard nested approach) and optimized polymerases and extension times, among other measures [56, 55]. They validated this approach by comparing linkage decay in a 50/50 mix of two laboratory strains. While no linkage decay was observed in this mixture, the reduced sequence similarity between the two strains may also have discouraged recombination [4, 5]. As further validation, they analyzed a natural control of an intra-host population founded by two distinct viruses. Despite the distinct founder viruses having high sequence similarity, nearly complete linkage was preserved after a single round of PCR amplification. Additionally, these control samples were from relatively high viral load time points (*≈* 10^5^ copies/mL) indicating that PCR recombination is extremely low even in the high viral load samples we analyzed. The agreement between the estimates from the Caskey et al. data [7] (produced via single genome sequencing) and the low viral load tertile of the Zanini et al. data also supports this point. While we cannot definitively say that PCR recombination did not in any way affect our results, these controls provide reassurance that rates of *in vitro* PCR recombination were extremely low.

It is important to note several additional caveats with our analysis: 1) Correlation does not imply causation. Although we report an association between recombination rate and viral load, it is possible that rapid recombination potentiates a higher viral load, and not that higher viral loads cause more coinfection and subsequent recombination. It is also possible that a third underlying factor causes both higher viral load and more frequent recombination. While we cannot directly disentangle causation in this study, the association still suggests that we must consider how recombination rates vary within and between individuals. 2) We analyzed a relatively small cohort of untreated participants, many of whom showed some degree of virologic control [56]. The same features that may permit these individuals to control the virus may also affect recombination processes within these viral populations. However, we note that individuals with low viral loads showed recombination rates consistent with other estimates in the literature [34, 6], while time points with less control exhibited elevated recombination rates more consistent with *in vitro* studies [27, 48]. 3) Most of our analysis derives from only a single data set [56] and we could not locate other data sets with the necessary characteristics (longitudinal, deeply-sequenced, abundant segregating sites, minimal PCR recombination, range of viral loads) for a full biological replicate. However, application to Caskey et al. [7] did produce a consistent estimate with the low viral load group. We are hopeful that as sequencing technologies proliferate, new data will be collected that permit further verification.

While RATS-LD is a broadly useful technique for quantifying rapid recombination from time series data, the sequencing strategy (especially the read length) limits the recombination rates that can be effectively inferred. Similar limitations have been noted in other work estimating linkage structure from pooled read data [14]. Low recombination rates may permit few or no recombination events to occur within the context of single reads. As a result, little breakdown of linkage will be observable and longer read sequencing data might be necessary to capture these events. As these sequencing approaches proliferate [15], we expect the utility of linkage-based techniques for quantifying recombination and other evolutionary forces to expand, even in populations with lower recombination rates. In the opposite direction, high recombination rates can result in extremely rapid decay of linkage which may saturate even between nearly adjacent loci if the sampling time points are insufficiently dense (see *ρ* = 10^*−*3^). This problem is potentially resolvable by denser sampling, although we note that the recombination rates explored here are some of the highest measured *in vivo* [40]. For the purpose of HIV, RATS-LD is well suited to the current state of short read paired-end next generation sequencing data.

Although we found that RATS-LD was appropriate for our uses in this paper, we offer a series of recommendations for assessing its appropriateness in other settings and the trustworthiness of its estimates. *1. Assess sampling characteristics:* we found RATS-LD performed best with at least 500 segregating sites and a sampling depth above 40 (Figures S11 & S8A). *2. Understand the* 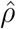 *performance range:* with paired end data spanning approximately 700 base pairs sampled on the order of 100 generations apart, we found that RATS-LD can discriminate well among recombination rates in the range of 5*×*10^*−*6^*−*5*×*10^*−*4^· Rates estimated outside of this range are generally untrustworthy under these sampling conditions. Longer range or temporally denser sequencing data may expand this accuracy range in the future. *3. Visualize linkage decay curves:* the raw data underlying these curves can be noisy, so we found binning by 500 bp *×* generation moving windows for visualization balanced stochasticity and signal. If these running average curves appear flat, RATS-LD will likely be unable to accurately estimate recombination rates. *4. Examine bootstraps:* bimodal bootstraps or those spanning multiple orders of magnitude reflect poor curve fitting and should be treated with caution.

More broadly, the biological phenomenon that we report here - association between higher recombination rate and higher population density - could be much more widespread in facultatively asexual populations where physical proximity can mediate the rate of recombination. Potentially relevant cases include viral homologous recombination (as shown here) and reassortment, but also rates of plasmid conjugation in bacteria, or sexual reproduction in hermaphroditic organisms or plants that do not self or outcross exclusively. Explicit attention to population density in mediating recombination rates stands to impact our understanding of species diversification in viruses and more broadly.

## Acknowledgements

The authors would like to thank Richard Neher for assistance accessing data from Participant 6, in addition to Kelley Harris, Benjamin Kerr, Daniel Reeves, Lillian Cohn, Elena Giorgi, Nandita Garud, the Cohn lab at Fred Hutchinson Cancer Center and the Feder lab at the University of Washington for useful discussions on this project and feedback on the manuscript. We will like to especially thank Fabio Zanini, two anonymous reviewers and the editor for providing valuable feedback on this work. EVR was supported by the UW Statistical Genetics Training Grant (T32GM081062). This independent research was supported by the Gilead Sciences Research Scholars Program in HIV.

## Materials & Methods

### Measuring Linkage Over Time

RATS-LD quantifies the linkage between pairs of segregating loci using D’ [28](1):

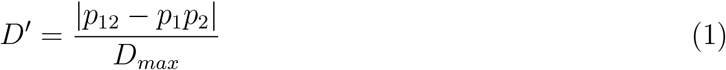

where *p*_1_ and *p*_2_ are the frequencies of the majority nucleotide at the corresponding locus, *p*_12_ is the frequency of the haplotype made up by the majority nucleotides at the two loci, and *D*_*max*_ is given by

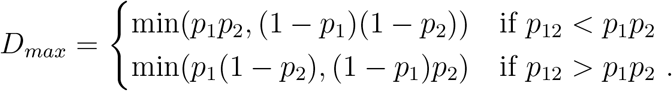

Estimation of *ρ* can also be performed using *D* rather than *D*′, but we found *D*′ was less affected by noise (Figure S16). Note, because linkage information is necessary to compute *D*′, pairs of SNPs must be close enough to be spanned by a sequencing read.

RATS-LD then computes the decay of linkage between those pairs of segregating loci at a later time point. For a given set of time points, *D*′ is expected to decay according to the following equation [18]:

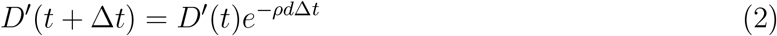

where *d* is the distance between the loci in nucleotides and Δ*t* is the time between the two time points in generations (assuming that *D*_*max*_ is similar over time). For the *in vivo* analysis, we converted the number of days between samples to generations by assuming an approximate rate of 1 generation per 2 days [39, 44]. We can rewrite (2) as

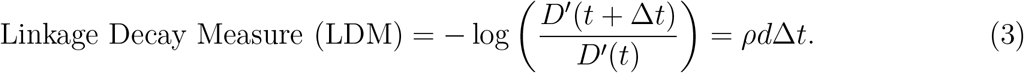

Both the left hand side of (3), which we will denote as the linkage decay measure, and *d*Δ*t* can be measured from the data directly, providing the opportunity to estimate *ρ*. However, the decay of linkage over time between two pairs of SNPs is a noisy measurement, so we fit this relationship over many pairs of SNPs and many time points, as described in the next section.

### Recombination Rate Estimation

After filtering the data as described below, we calculated linkage decay measures (LDMs) (3) for each pair of segregating loci at each pair of time points. Then we estimated recombination rates by fitting the decay of these LDMs over the distance between loci and time between samples (*d*Δ*t*). Note: we found that with sampling schemes in the simulated and *in vivo* data sets, HIV’s rapid turnover led to the linkage decay signal being predominated by decay over distance rather than time.

Equation (3) suggests a linear relationship between the linkage decay measure and *d*Δ*t* with slope *ρ*. However, two factors complicate this expected pattern: 1) sampling error makes it difficult to estimate small differences between *D*′ values accurately. As a result, a decrease of *D*′ from 0.01 to 0.001 may be driven entirely by sampling error, whereas a decrease from 0.5 to 0.05 likely reflects true decay in linkage, although they result in equal LDMs. At large values of *d*Δ*t, D*′ values are expected to become increasingly small, so the effect of sampling error becomes pronounced.

2) Over long periods of time, other evolutionary forces - selection, recurrent mutation, population turnover - are more likely to affect linkage relationships between two loci. Therefore, to overcome these challenges, we fit the functional form

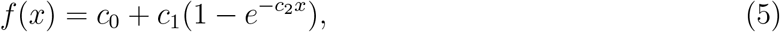

where 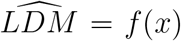 and *x* = *d*Δ*t*. The slope of the curve as *d*Δ*→* 0 should still be *ρ* since, in this range, there is minimal time for evolutionary processes to act and tight linkage leads to large *D*′ values which are not as affected by sampling noise as detailed above. This yields a final estimate, given by the derivative of (5) at *d*Δ*t* = 0 which is

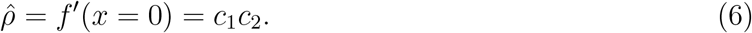

Thus, RATS-LD quantifies the recombination rate by fitting the relationship between linkage decay and time-scaled distance (*d*Δ*t*) then calculating the derivative of the fit near *d*Δ*t* = 0.

### Curve Fitting

For all analyses except those shown in Figures S2 & S3, we fit the curve form (5) to the LDM data via non-linear least squares [10]. Prior to performing the fits, we restricted the data to pairs of SNPs with *d*Δ*t <* 5*×*10^4^·bp generations to boost RATS-LD’s performance in the range of previous HIV recombination rate estimates (Figure S11A).

To confirm that the slope of the linkage decay curve near *d*Δ*t* = 0 does capture the true recombination rate, we performed linear fits to the LDM data which are shown in Figures S2 & S3. Since the linear fits are only accurate when considering LDMs prior to saturation, we performed these fits to data only including LDMs with *d*Δ*t* < 500, 1000, 2000, 10000, or 50000 bp · generations so that at least one *d*Δ*t* fit threshold would capture the pre-saturation behavior for each value of *ρ* in the simulated data.

We fit these relationships with linear regression [10] and the slope of the fit was the estimated 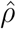.

### Bootstrapped Confidence Intervals

To create bootstrapped confidence intervals for RATS-LD’s recombination rate estimates, we sampled the segregating loci with replacement and repeated estimation using only the LDMs where both loci were sampled. Each bootstrap sample included the same number of segregating loci as was initially used for estimation. The example fits displayed for the bootstrapped confidence intervals use the set of coefficients that provided the estimate at the given quantile (2.5%, 50%, 97.5%) of the 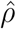 distribution (rather than the quantiles of the coefficient distributions themselves). The primary 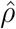 reported for each analysis is the median of the bootstrapped distribution.

### Simulated Data

To validate RATS-LD’s ability to quantify recombination rates, we simulated populations with known recombination rates using SLiM [17]. We performed RATS-LD’s initial validation on a simulated genomic region of length 700bp, with a mutation rate of *μ*= 10^*−*6^ mutations/bp/generation [2, 1, 46] and effective population size of *N*_*e*_ = 10^4^ [2, 11]. The simulated recombination rates *ρ* ranged from 2 *×*10^*−*6^ to 10^*−*3^ recombinations/bp/generation to broadly encapsulate the range of previously estimated HIV recombination rates [6, 34, 51]. Note: for the purpose of the discrimination analysis only (Figure 2D,E), we also ran neutral simulations in an expanded range of *ρ* = 10^*−*6^ to *ρ* = 5*×*10^*−*3^. Each simulation was run for 5*×*10^4^ generations before samples from five evenly spaced time points were collected at intervals of 150 generations. Each sample contained *M* = 800 genomes to approximately match the median number of templates in each of the *in vivo* sequencing reactions [56]. RATS-LD requires many pairs of SNPs across multiple time points to fit Equation 4. Therefore, data pooling across simulations is required for estimation. Since our power analysis indicated that estimation is most accurate when there are *>* 500 loci available for estimation (Fig. S11C), for each recombination rate estimate we grouped 5 simulations and sampled 200 segregating loci from each simulation to form groups with *≈*1000 segregating loci (Table S1).

To assess RATS-LD’s performance in the presence of selection, we performed additional simulations with selection coefficients designed to mimic intra-host HIV evolution more closely. These selection simulations included a genomic region of length 700bp, mutation rate *μ*= 10^*−*5^ mutations/bp/generation [2, 1, 46], and effective population size *N*_*e*_ = 10^4^ [2, 11]. Selection coefficients were drawn from the following distribution of fitness effects: a small number of mutations (*≈*0.1%) were strongly beneficial with *s* = 0.05. Most mutations (*≈*70%) were non-synonymous mutations [24] and were strongly deleterious with selection coefficient *s* =*−*0.1 [57]. The remaining *≈*30% of mutations represented synonymous mutations and were neutral with *s* = 0 (frequency *≈*10%) or slightly deleterious with *s* =*−*0.01 (frequency *≈*20%) [57]. Each simulation was run for 50 generations before 9 evenly spaced samples of *M* = 800 templates each were collected at intervals of 150 generations. The number of sampling time points was increased to 9 so that the data sets would encompass the maximum length of sampling for participants in the *in vivo* data set [56]. Under these simulated conditions, the proportion of segregating sites per time point pair in the selection simulations matched the *in vivo* data much more closely than the neutral simulations (Figure S17). For each estimate, we grouped 50 simulation runs which provided 700 to 2000 segregating loci to ensure that RATS-LD was well-powered (Table S1). Note: due to the simulations’ shorter genomic length (700 bp) vs. the average *in vivo* fragment length (mean *≈*1800 bp), a group of 10 simulations is approximately equivalent to the data from one participant. The selection simulations described in this paragraph are referred to as the ‘baseline selection simulations’.

We also performed selection simulations to test for impacts of other population factors on RATS-LD estimates. First, we tested the impact of population substructure by simulating populations consisting of two separate sub-populations at *ρ* = 10^*−*4^ and *ρ* = 10^*−*5^. Viruses migrated between the two sub-populations with backward migration rates of *m* = 10^*−*5^, 10^*−*4^, 10^*−*3^, and 10^*−*2^. The extremes of these values were drawn from the literature [29, 12]. Each sub-population had size 5 *×*10^3^ resulting in full population sizes of *N*_*e*_ = 10^4^ (consistent with the baseline selection simulations). Both sub-populations contributed equally to the sampled viruses. All other parameters in the substructure simulations matched those in the baseline selection simulations. Second, we tested the impact of varying effective population size. These simulations had the same parameters as the baseline selection simulations except that we simulated a wider range of effective population sizes (*N*_*e*_ = *×*10^3^, 10^4^, and 5*×*10^4^) for a subset of recombination rates (*ρ* = 2*×* 10^*−*6^, 10^*−*5^, 2*×*10^*−*5^, 10^*−*4^, *×*10^*−*4^, and 10^*−*3^). Third, we tested the impact of varying sampling depths. These simulations have the same parameters as the baseline selection simulations, but we varied the sampling depth of *M* to be 10, 20, 40, and 80. At lower sampling depths, many more simulations (up to 5000 per estimation at *M* = 10) were required to form groups of 500*−*1000 segregating loci. Since larger numbers of independent observations reduce noise and improve RATS-LD accuracy, these estimates represent best case scenarios for each sampling depth.

To match the sequencing coverage in the *in vivo* data, we created a procedure to downsample the simulations to mimic the pairwise coverage of two loci *d* nucleotides apart. To do this, we created an empirical distribution of reads overlapping a locus at an arbitrary position 0 (Figure S18A). First, we simulated the expected positions of 10^5^ reads with start positions uniformly distributed between genomic positions -700 and 0 (in bp). For each start position, we uniformly sampled a total end to end read length between 400 and 700 bp, which was the observed read length distribution in Zanini et al., 2015. In addition, we only considered 300 bp of coverage extending inwards from each end of the read, to match the paired end sequencing study design [56, 55]. As a result, longer reads did not have coverage in their centers. We discarded any simulated reads that did not capture genomic position 0. At this point, we formed the empirical distribution by computing the number of reads that overlapped position 0 and a position *d* nucleotides away in either direction. The resulting empirical distribution had high coverage for nearby loci and decreasing coverage as *d* increased (Figure S18C). This paired-end read sampling technique mimicked the *in vivo* sampling scheme and provided coverage similar to the *in vivo* data ([56]) (Figure S18B). Then, for each pair of SNPs in the SLiM data at a distance of *d* nucleotides apart, we downsampled their coverage to match the empirical number of overlapping reads for two SNPs at a distance of *d* nucleotides. This procedure was used to downsample coverage for all simulation scenarios described.

### *In vivo* Data

We used longitudinal HIV deep-sequencing data from Zanini et al. 2015 for *in vivo* recombination rate estimation [55, 56]. The samples were collected from the blood plasma of 11 participants who were diagnosed with HIV-1 in Sweden from 1990 – 2003 and were not on antiretroviral treatment. Six to twelve time points are included per participant with both sequencing and viral load information at most time points. The sequences span the full HIV genome and were generated using 6 overlapping amplicons, but we excluded fragment 5 (which contains *env*) in our analysis due to its lower rates of template recovery. The sequencing reactions used paired-end reads amplified via a protocol designed to minimize PCR recombination (see Discussion). In particular, the protocol used only a single round of PCR (rather than a nested approach). We used arrays of co-occurrences of alleles across pairs of loci (available at https://hiv.biozentrum.unibas.ch), which had undergone Zanini et al.’s quality assurance process for read mapping and filtering. This process included rigorous checks for base quality, mapping errors, and sample contamination. We also used the estimated dates of infection (EDI) as provided by Zanini et al., and each time point was measured relative to the EDI for the corresponding participant.

To assess if higher viral load is associated with a difference in recombination rate, we sorted pairs of time points into groups based on their average viral load. We started by matching genotype information from each sampling time point with a viral load measurement near the same time. For the majority of the data, the date of sample collection exactly matched the date of the viral load measurement. However, some samples from participants 4 and 7 did not have a concurrent viral load measurement and were therefore matched to viral load measurements within 100 days of the sample’s collection. Additionally, data from Participant 10 was omitted from the analysis due to nearly all of the time points lacking viral load measurements within 100 days. For each pair of time points, we calculated the average viral load by taking the mean of the viral load measurements matched to those two samples. For pairs of time points with intervening viral load measurements, we required that those viral loads stayed within the bounds of the viral loads measured at the starting and ending time points. Pairs of time points with an intervening measurement that violated these boundaries were excluded from the analysis to ensure that the average viral loads used to group the data are as accurate as possible.

For the quantile analyses, the LDMs were labeled with the average viral loads across their corresponding time point pairs and divided based on average load to make two (Figures 5 & S15) or three (Figures 4 & S12) groups with approximately equal numbers of LDMs (Table S1). Since a single pair of time points could have many corresponding LDMs, the size of the groups sometimes varied by *≈*1000 measurements since all LDMs per time point pair were included in the same group. For the threshold analysis shown in Figures S13 and S14, the pairs of time points were placed into groups based on whether their average viral load was above or below a given threshold, and we then applied RATS-LD separately to the high and low viral load groups.

We also performed three additional checks to confirm our observations. First, to ensure the pattern observed in Figure 4C was not solely driven by data from a single participant, we performed a leave-one-out analysis in which we repeated estimation excluding all data from one participant at a time. Second, to ensure that our results were not driven by two individuals who experienced superinfection, we performed an analysis leaving out both of these individuals simultaneously. Third, we analyzed a data set of viral sequences described in Caskey et al. 2017, which served as a partial biological replicate [7]. These sequences were collected from 15 study participants with rebounding viral populations after a single infusion of a broadly neutralizing antibody treatment. All participants were either ART naive or were off ART for 8 weeks prior to the study start. We used only the data produced via single genome sequencing, which provided an average sequencing depth of *≈*23 reads. We aligned the sequences using the LANL Codon Alignment v2.1.0 tool (https://www.hiv.lanl.gov) and manually checked alignments before proceeding with the estimation process. Additionally, we used only the data starting at 4 weeks post treatment, when viral populations were rebounding. Prior to the estimation, we grouped the full data set due to the low sequencing depth and relatively low viral loads (Figure S8B).

### Data Filtering

We started with data from all time points and all pairs of SNPs which were close enough to be spanned by sequencing reads. Then, we used several filtering steps in both the simulated and *in vivo* data sets to reduce noise. 1) We required that at least 10 reads span a pair of loci for that pair to be included in the analysis. 2) SNPs at each time point were included only if the minor allele had a frequency *>* 1% at that time point. We chose this cutoff since Zanini et al., 2015 report that SNPs’ frequencies can be estimated down to 1% accuracy in the *in vivo* data set we analyzed [56, 55]. Any SNPs transiently under 1% frequency were not included in the analysis at the corresponding time point, but were included at other time points where they reached the frequency threshold. In the Caskey et al. dataset, we used a *>* 10% allele frequency cutoff due to its lower sampling depth. For the linkage analysis displayed in Figure S6, we required a minor allele frequency *>* 20% to replicate the filtering conditions used to generate Figure 7 of Zanini et al., 2015 [56]. Filtering step 2 prevented false linkage signals from very low frequency alleles appearing and disappearing between time points. 3) All four possible haplotype combinations of the major and minor alleles at a pair of loci had to be observed at a given time point to be included in the analysis at that time point. Note: for the analysis with the *D* statistic (Figure S16), we did not require that all four possible haplotypes were present. 4) Because the investigation focused on the decay of linkage across pairs of time points, we required that *D*′ *>* 0.2 at the first time point in each pair (or *D >* 0.0075 for the analysis using D) since decay in larger *D*′ values is less likely to be a function of noise. RATS-LD’s ability to discriminate between underlying recombination rates is robust to changes in this threshold. 5) Lastly, we excluded time point pairs that spanned less than 50 days, since the signal of linkage decay at such short timescales is also very susceptible to noise. For counts of segregating loci and their corresponding LDMs which passed these filtering steps, consult supplemental tables S1 and S2.

### Discrimination Analysis

To validate that RATS-LD can discriminate between different recombination rates, we randomly paired simulations with recombination rates that were a factor of 1, 1.1, 1.2, 1.2, 2, or 5 apart. For each value of *ρ* we used 20 data sets of *≈*1, 000 neutrally evolving loci which were simulated in SLiM as described above. We applied RATS-LD to each simulated data set to produce a 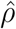 estimate given by the median of its corresponding bootstrapped distribution (1000 bootstraps). To compare two given *ρ* values, we randomly paired these estimates to create 20 matched pairs. For each matched pair with true recombination rates *ρ*_1_ and *ρ*_2_, we computed two comparison metrics. 1) Simple rank order: we determined whether the individual estimates 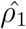 and 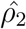were correctly ordered such that 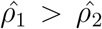 if *ρ*_1_ *> ρ*_2_. If *ρ*_1_ = *ρ*_2_, we checked agreement against an arbitrary ordering. 2) Bootstrapped significance: we determined whether the bootstrapped 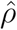 distributions from the data sets overlapped. We reshuffled the pairings and recomputed the two metrics 500 times.

## Data availability

The primary data set used in this study (viral loads and paired participant read counts from [56]) can be downloaded from https://hiv.biozentrum.unibas.ch/. The single genome sequencing data from [7] can be downloaded from genbank (accession numbers KY323724–KY324834) while the viral load measurements from the study are listed in Supplementary Table 3 of Caskey et al., 2017.

### Code availability

All code to reproduce these analyses can be found at

https://github.com/federlab/hiv-recombination.

---

**Table S1:** Counts of segregating loci and linkage decay measures per estimation group. The recorded number of segregating sites includes all unique loci segregating at any time point in the estimation group (i.e., upper half or lower half). As a result, *in vivo* loci can be counted in multiple groups if they segregate in both those groups. The mean and standard deviation of counts are recorded for simulated data.

| Estimation Group | # of Segregating Sites | # of LDMs |
| --- | --- | --- |
| Neutral Simulations | 996 ± 9 | 66,465 ± 13,591 |
| Selection Simulations | 1,812 ± 798 | 3,993 ± 1,793 |
| Lower Tertile (Fig. 4 Data) | 3,285 | 25,579 |
| Middle Tertile (Fig. 4 Data) | 2,530 | 19,353 |
| Upper Tertile (Fig. 4 Data) | 2,306 | 23,778 |
| Upper Half (Fig. 5 Data Par.1) | 758 | 8,708 |
| Lower Half (Fig. 5 Data Par.1) | 570 | 5,111 |
| Upper Half (Fig. S15 Data Par.3) | 670 | 5,196 |
| Lower Half (Fig. S15 Data Par.3) | 558 | 4,594 |
| Upper Half (Fig. S15 Data Par.4) | 556 | 5,661 |
| Lower Half (Fig. S15 Data Par.4) | 525 | 3,119 |
| Upper Half (Fig. S15 Data Par.7) | 996 | 14,800 |
| Lower Half (Fig. S15 Data Par.7) | 876 | 10,045 |

**Table S2:** Counts of segregating loci and linkage decay measures per participant. The recorded number of segregating sites includes all unique loci that were segregating at any time point.

|  | # of Segregating Sites | # of LDMs |
| --- | --- | --- |
| Participant 1 | 823 | 12,194 |
| Participant 2 | 308 | 1,335 |
| Participant 3 | 768 | 8,492 |
| Participant 4 | 663 | 8,217 |
| Participant 5 | 400 | 3,433 |
| Participant 6 | 172 | 546 |
| Participant 7 | 1,096 | 23,577 |
| Participant 8 | 348 | 1,943 |
| Participant 9 | 422 | 4,197 |
| Participant 11 | 500 | 4,775 |

**Figure S1:**
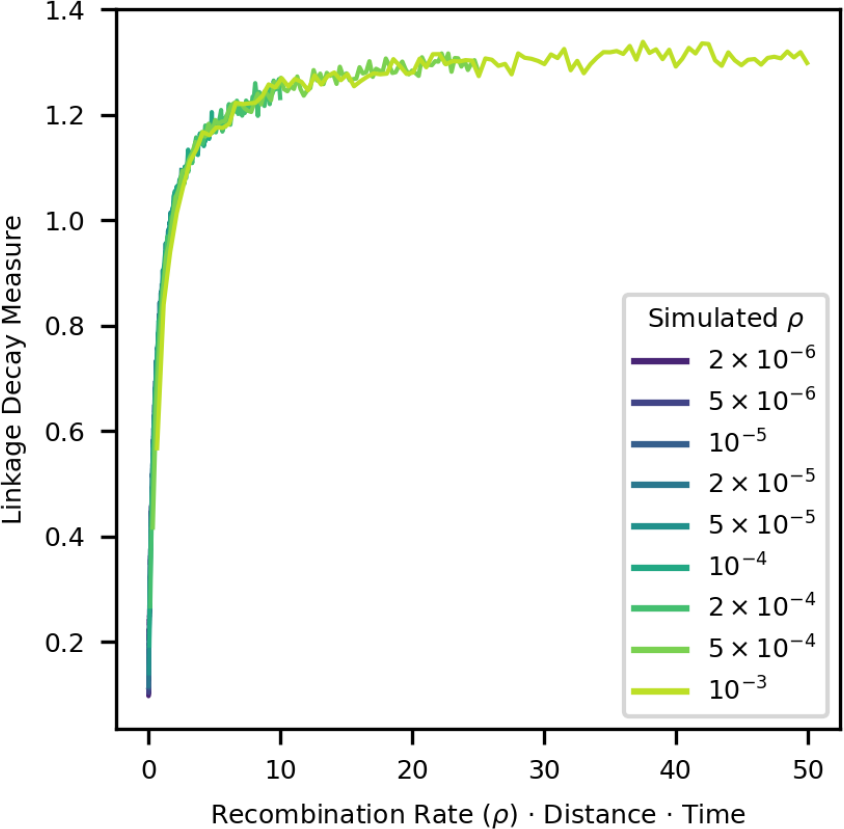
Average linkage decay measure as a function of recombination rate ·distance ·time reveals similar decay patterns across simulations with varying *ρ*. Binned average LDMs across the neutrally simulated data sets are shown (window size = 500 bp · generations). As expected, these curves collapse on top of each other when the known recombination rate *ρ* from the simulation is also used as a factor on the x-axis, verifying that the varying saturation patterns in Figure 2A result from recombination rate variation.

**Figure S2:**
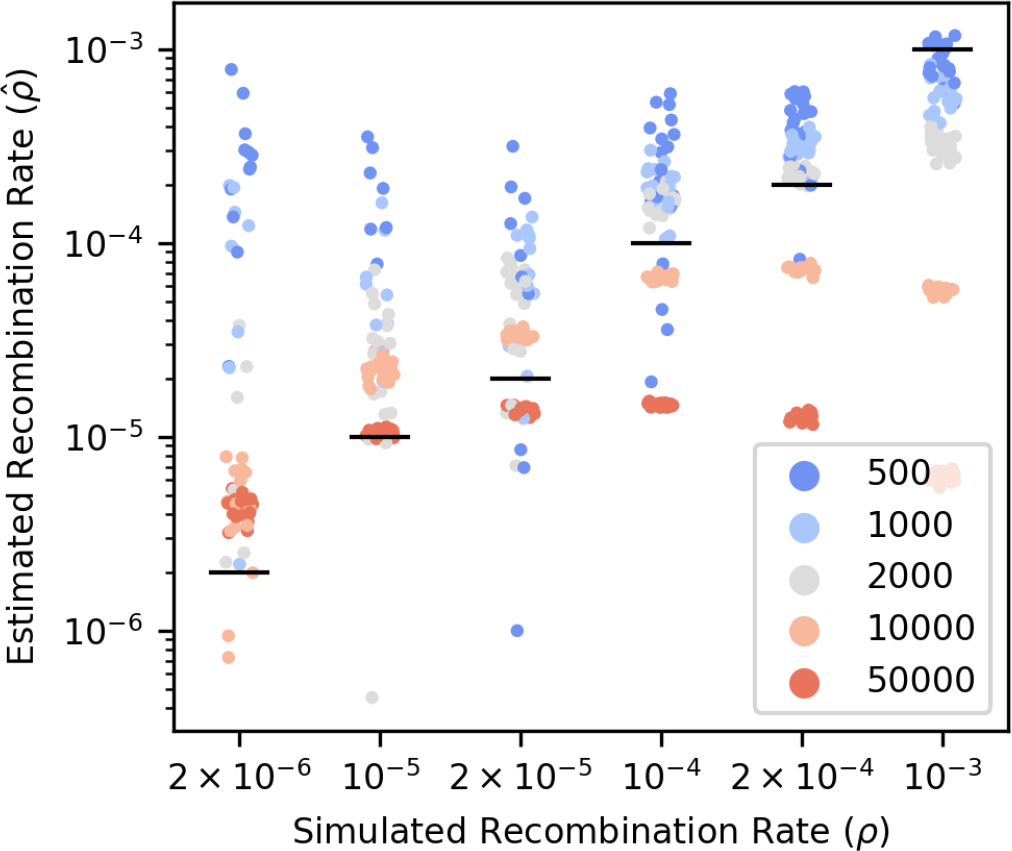
Fitting lines to linkage decay measures with increasing lengths of distance · time (*d*Δ*t*) shows that recombination rates are best recaptured at low *d*Δ*t* for high recombination rates while the opposite is true for low recombination rates. Note that 77/600 fits resulted in negative estimates which are not shown on this plot due to the necessity for a log scale y-axis. All of the negative estimates were from fits with *d*Δ*t* =*<* 2000. Examples of the fits used to generate this figure are shown in Figure S3. Each point corresponds to the estimate produced from a single line fit.

**Figure S3:**
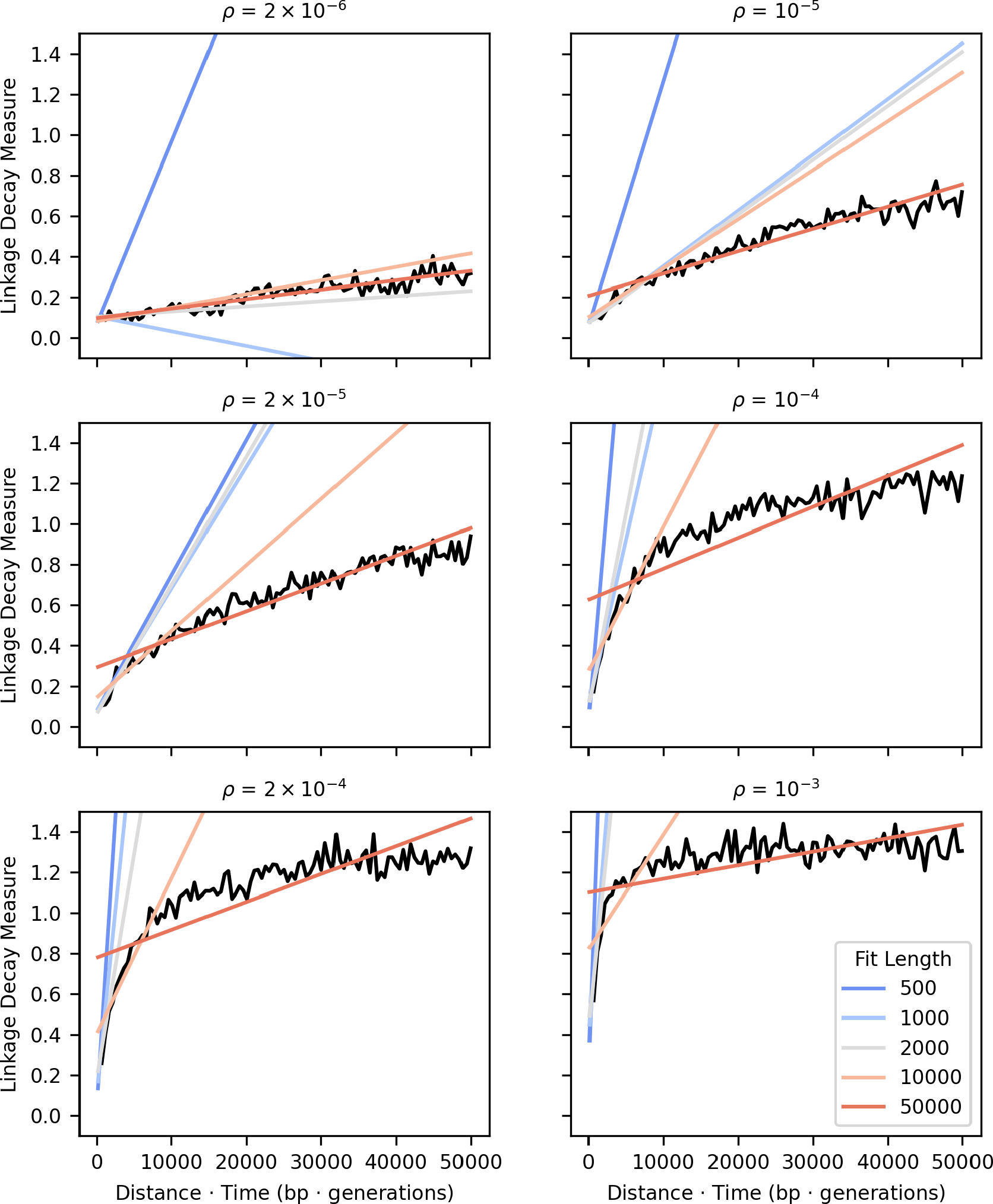
Example line fits to data truncated at increasing distance ·time values capture differing curve behaviors depending on the underlying recombination rate *ρ*. The black line in each panel represents a binned average of the linkage decay measures for the panel’s corresponding value of *ρ* (moving average for display purposes only, window size = 500 bp · generations). Linear fits to the LDM data truncated at varying values (shown in different colors), are plotted on top of the binned average.

**Figure S4:**
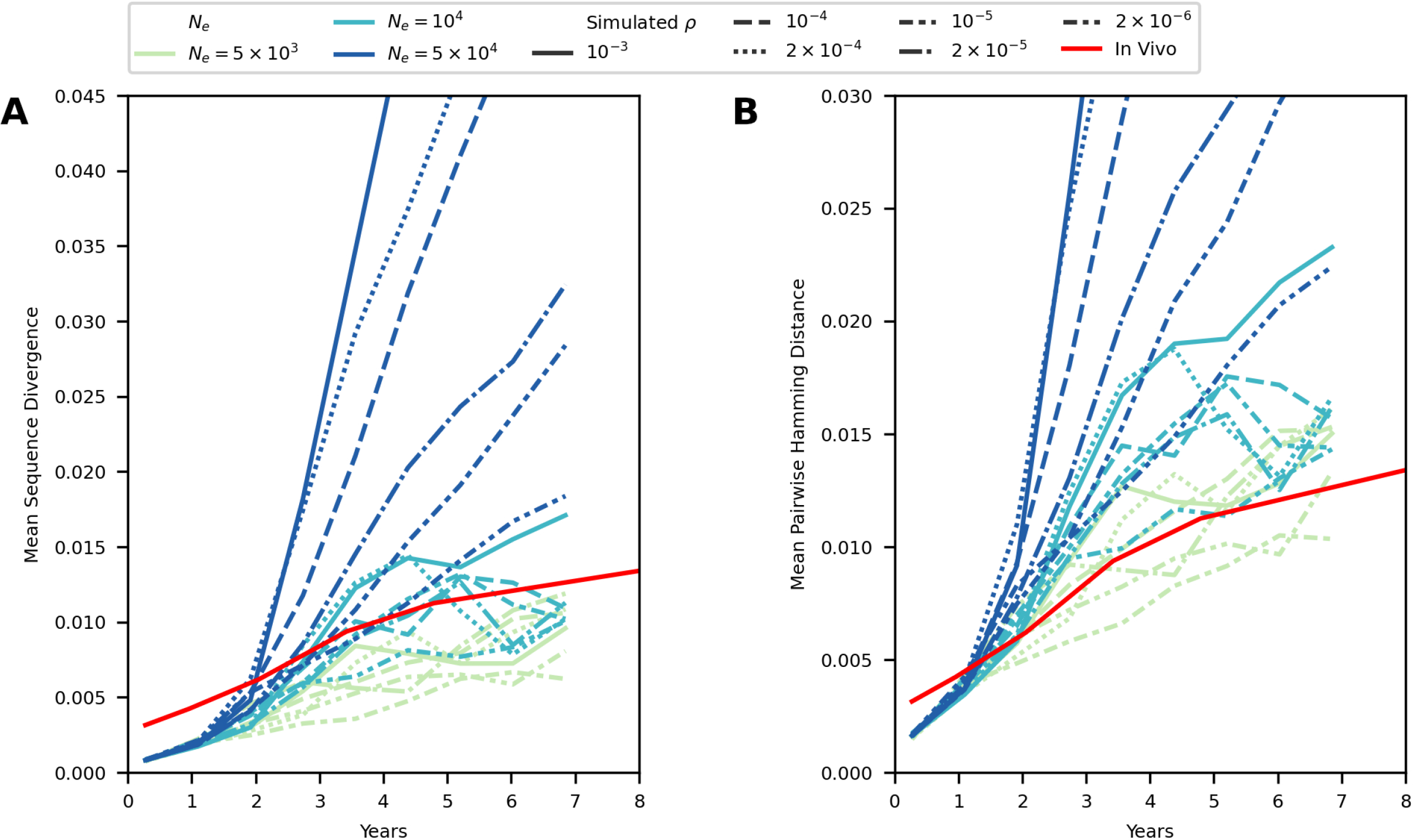
Selection simulations approximately match sequence diversity and divergence observed *in vivo*. Mean sequence divergence **(A)** and mean pairwise Hamming distance **(B)**are plotted over time. Selection simulations were performed with effective population sizes of 5*×*10^3^, 10^4^, or 5*×*10^4^. Each line represents data from 500 simulations and its line type corresponds with the underlying *ρ* value. *In vivo* data from Zanini et al. 2015 (Figure 5, panels A and B [56]) are plotted in red for comparison, averaged across all proteins and mutation types.

**Figure S5:**
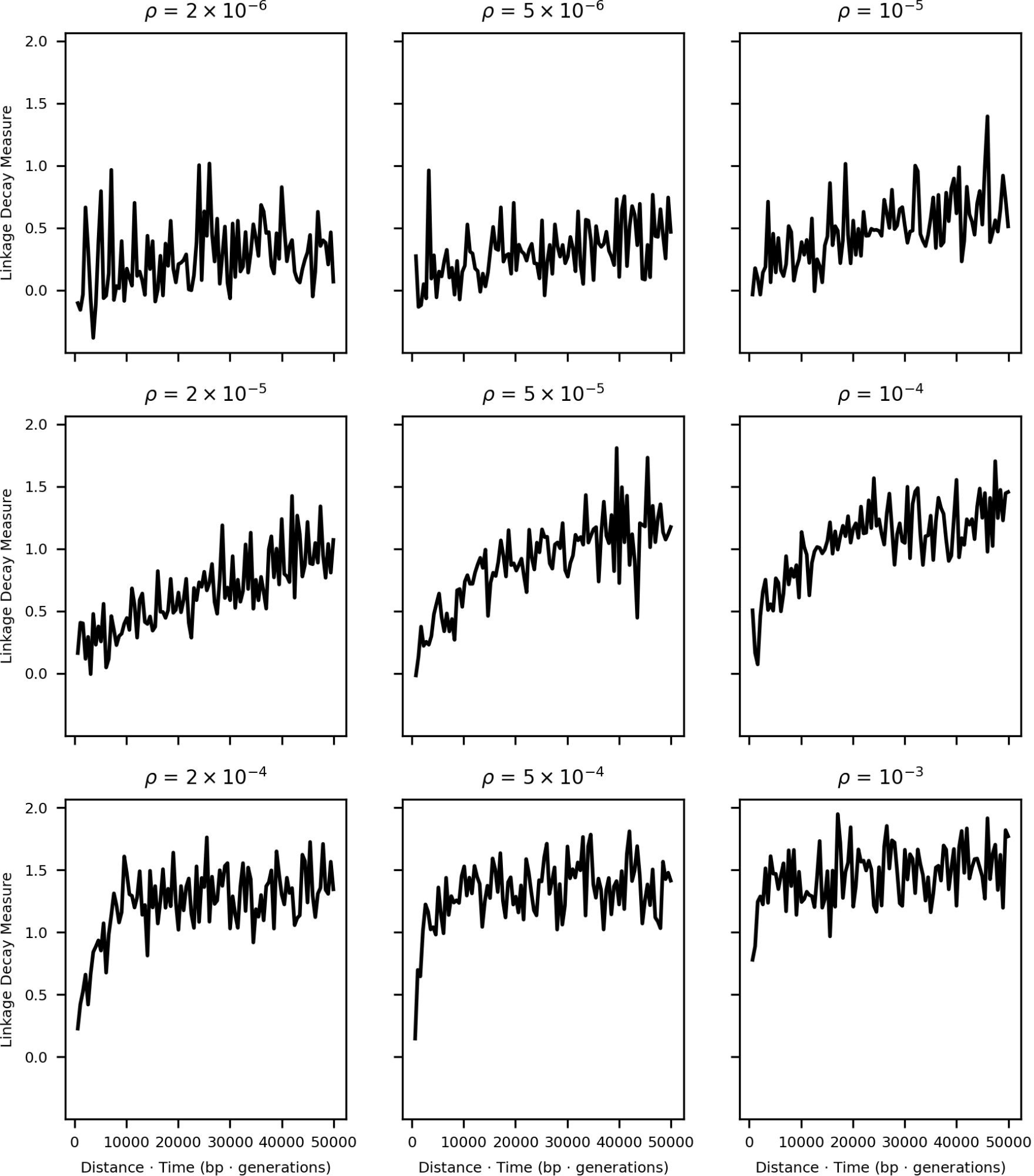
Example LDM curves for the simulated selection data set. A single example linkage decay curve is plotted for each simulated *ρ* value in the selection data set. Each curve represents the binned average LDM (window size = 500bp) from one estimation group. For direct comparisons to the neutral and *in vivo* data sets, see Figures S3 and 4B, respectively.

**Figure S6:**
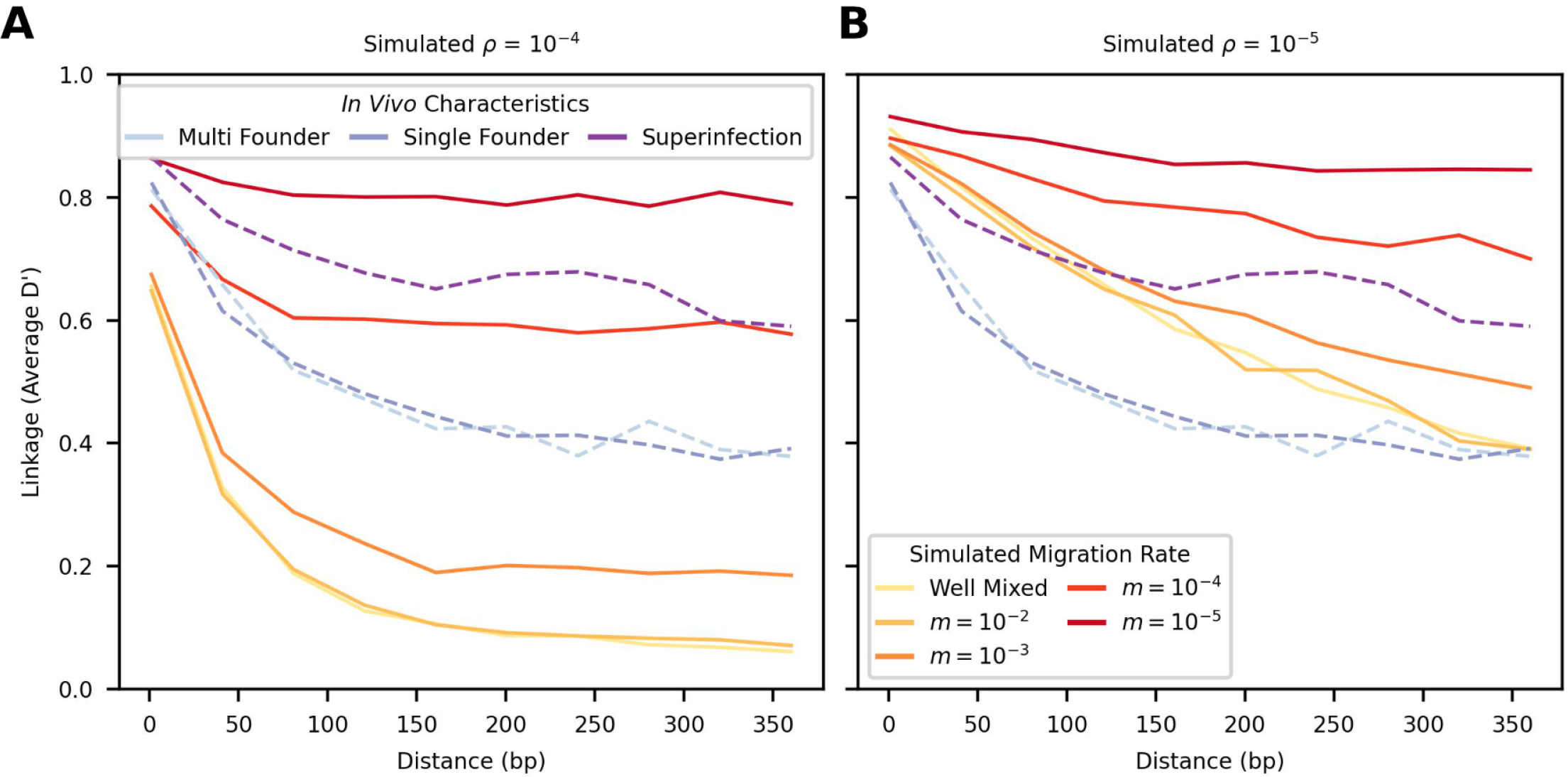
Simulated scenarios of population substructure encapsulate linkage decay patterns observed *in vivo*. The average D’ value in simulated data sets sampled from two sub-populations interconnected by migration rate *m* is plotted at different genetic distances for *ρ* = 10^*−*4^ **(A)** and *ρ* = 10^*−*5^ **(B)**. In each panel, the ‘well-mixed’ category corresponds to simulated data sets with no population substructure. Each simulated line was formed by taking a binned average (window size = 40bp) of D’ across 250 data sets (see Materials & Methods for full simulation and sampling details). The *in vivo* linkage decay is plotted with identical dashed lines on top of both panels for reference. These curves are colored depending on whether the intrahost population had diversity consistent with superinfection (Participants 4 and 7), transmission of multiple closely related founder viruses from the same individual (Participants 3 and 10), or transmission of a single founder virus (all other participants) [56].

**Figure S7:**
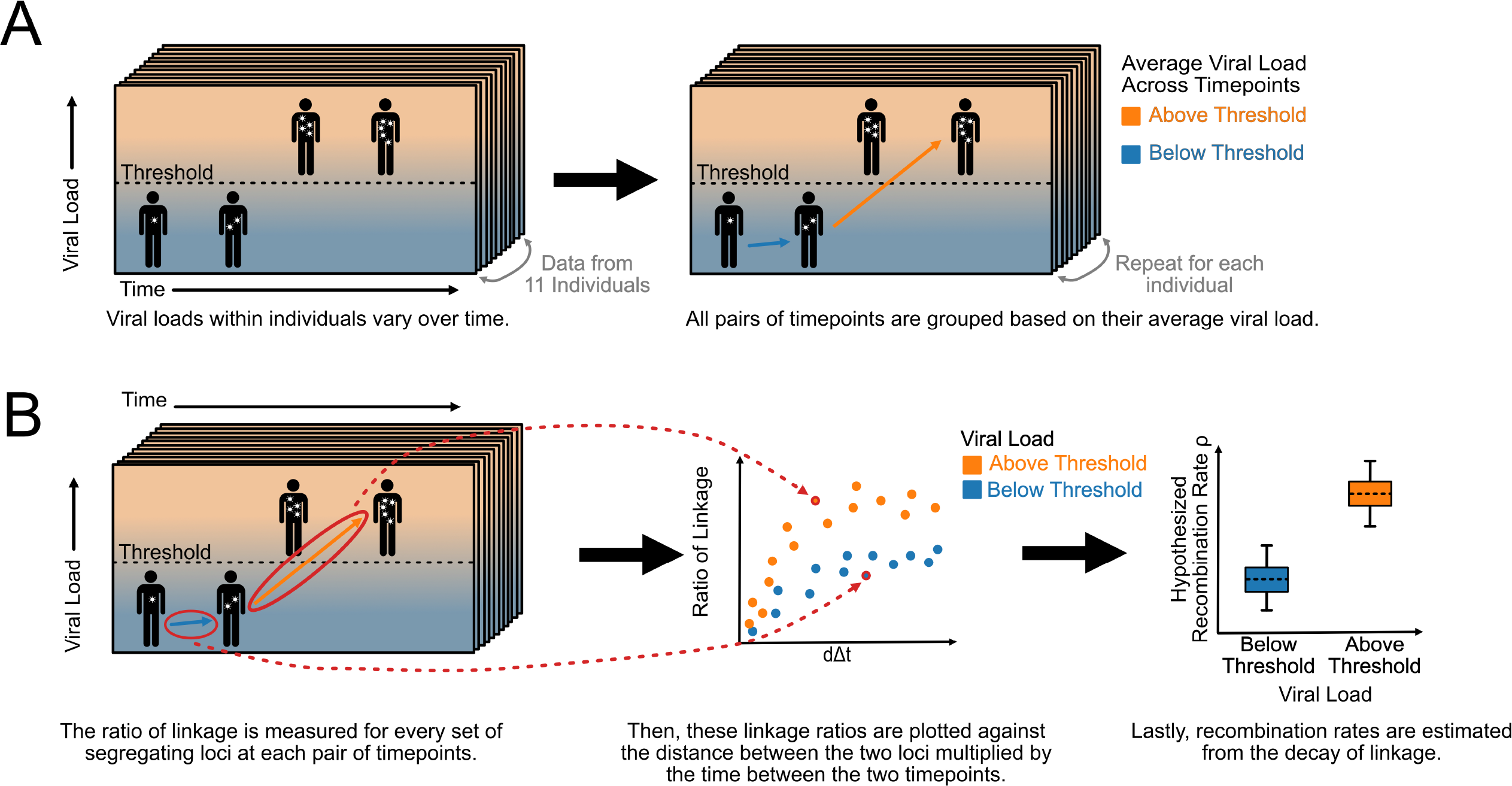
Workflow for *in vivo* viral load analysis.

**Figure S8:**
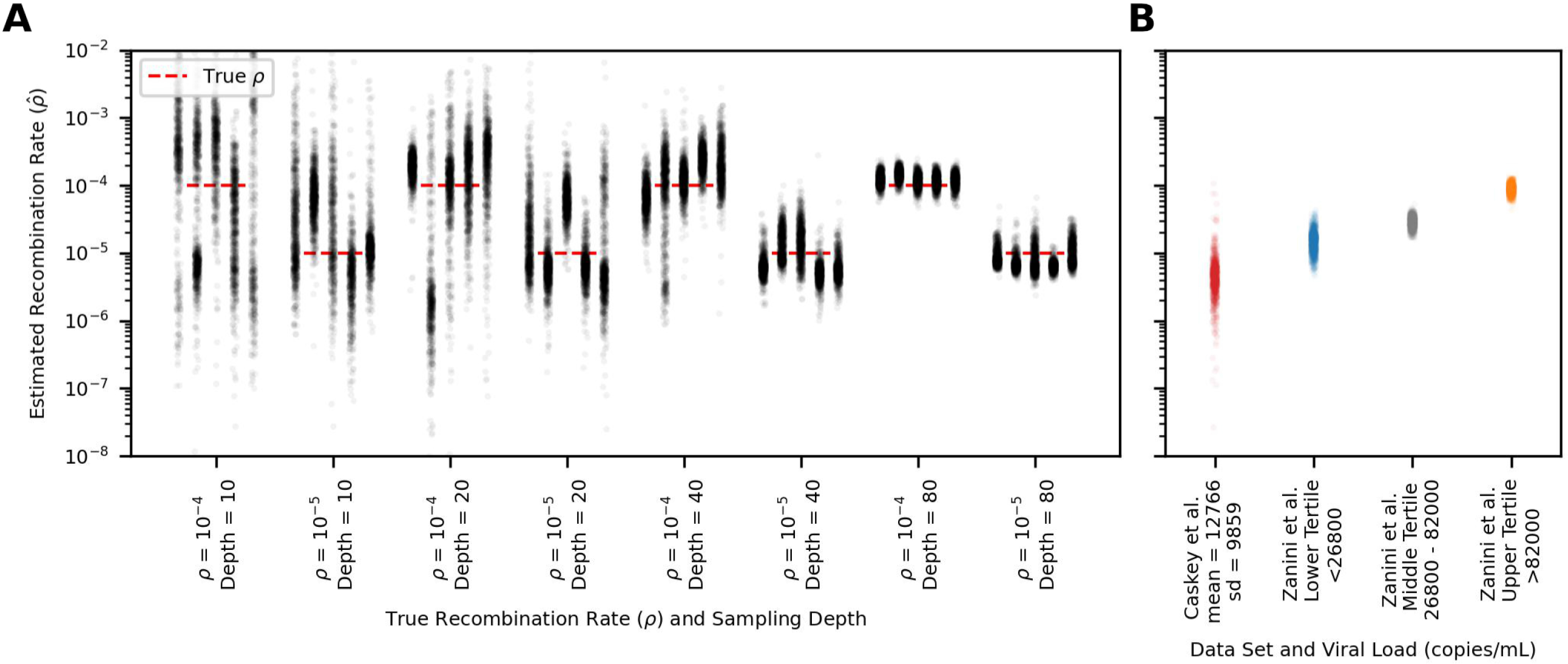
Low sampling depth is associated with wide *ρ* bootstraps, but *in vivo* data sets have sufficient depth for RATS-LD estimation. **(A)** Five estimation groups with 1000 bootstrapped 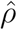 values are plotted for multiple sampling depths (10, 20, 40, or 80) and *ρ* values (10^*−*5^ or 10^*−*4^). **(B)** 1000 bootstraps are plotted for each of the viral load tertiles in the Zanini et al. 2015 dataset [56] alongside a single estimation group for the Caskey et al., 2017 [7] single genome sequencing data set (See Materials & Methods for more details). The cutoff viral load is indicated in the x-axis labels for the Zanini et al. data while the Caskey et al. data set is labeled with the mean and standard deviation of the viral loads associated with LDMs used in its analysis. The variation in the Caskey bootstraps is consistent with variation in the simulated data with approximately the same sampling depth (*≈* 23).

**Figure S9:**
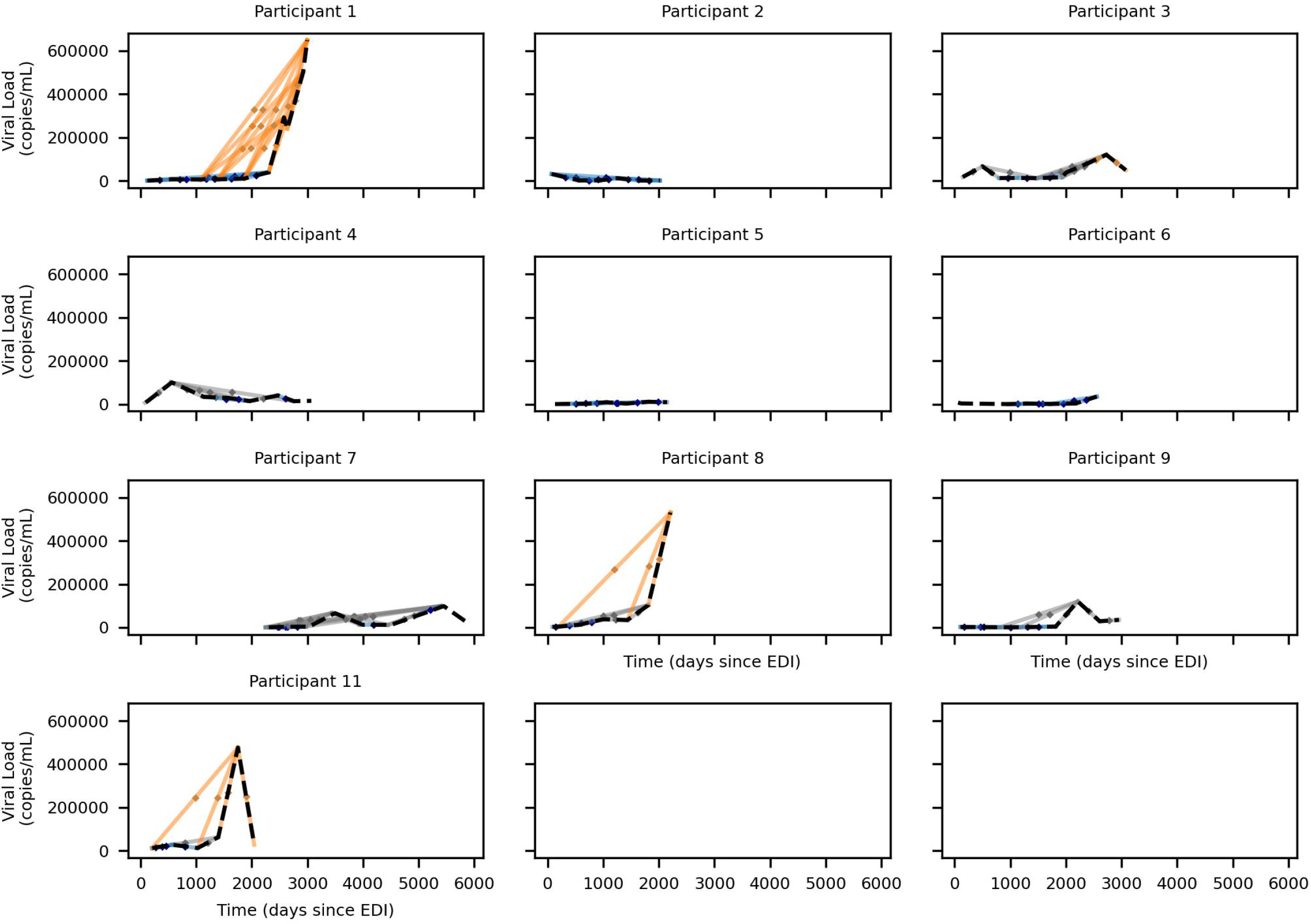
Viral load trajectories of individuals over time. In each plot, the black dashed line indicates the viral load measurements over time for a given individual. Each solid line segment connects a pair of time points included in the analysis and the time point pair’s average viral load is indicated by a dot at the center of the segment. The segments are colored blue (lower tertile *<* 26, 800), gray (middle tertile 26, 800*−*82, 000), or orange (upper tertile *>* 82, 000) based on their viral load groups.

**Figure S10:**
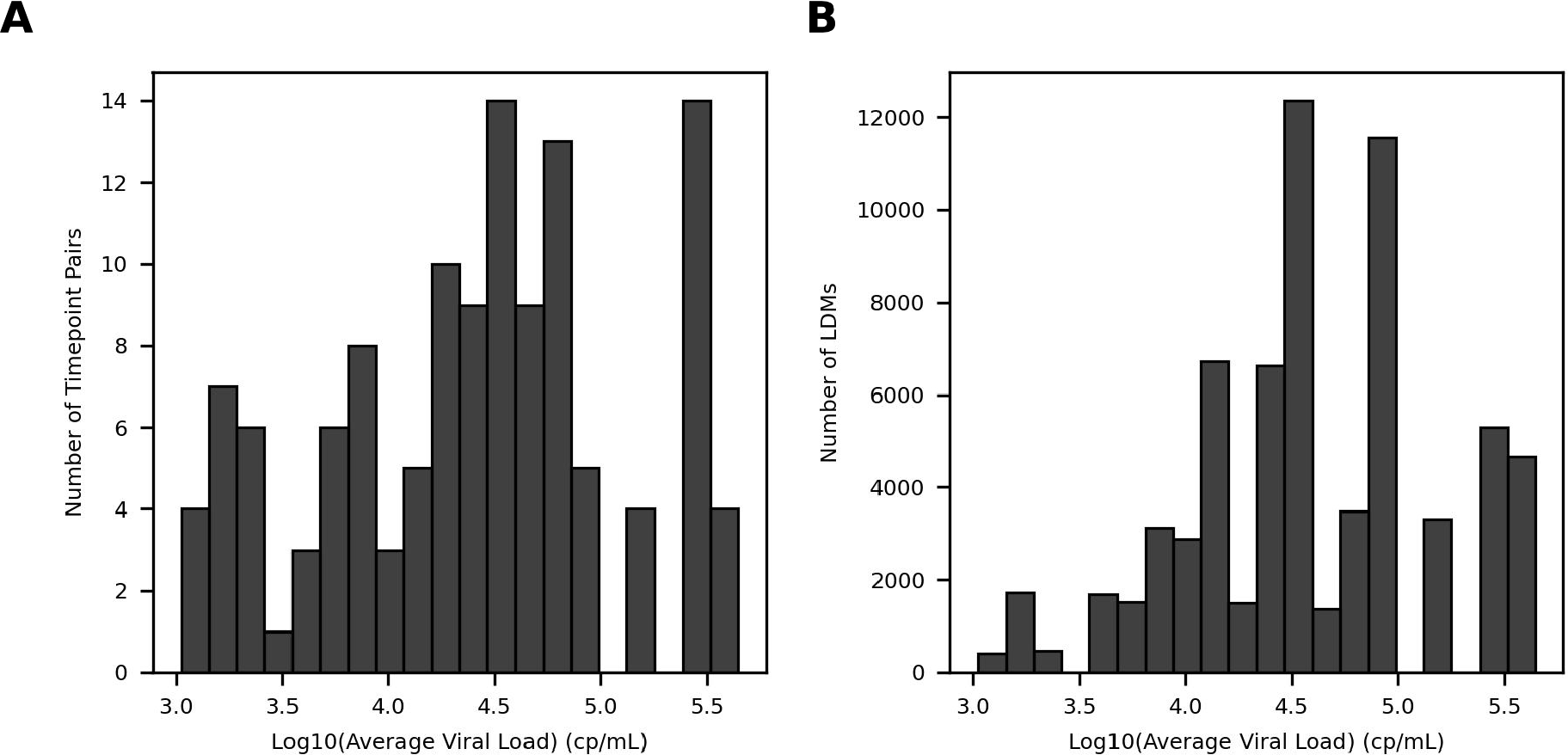
Distributions of average time point pair viral loads in Zanini et al. [56]. Histogram of viral load measurements averaged over each time point pair used in the analysis (Figure 4) binned by **(A)** number of timepoints and **(B)** number of LDMs.

**Figure S11:**
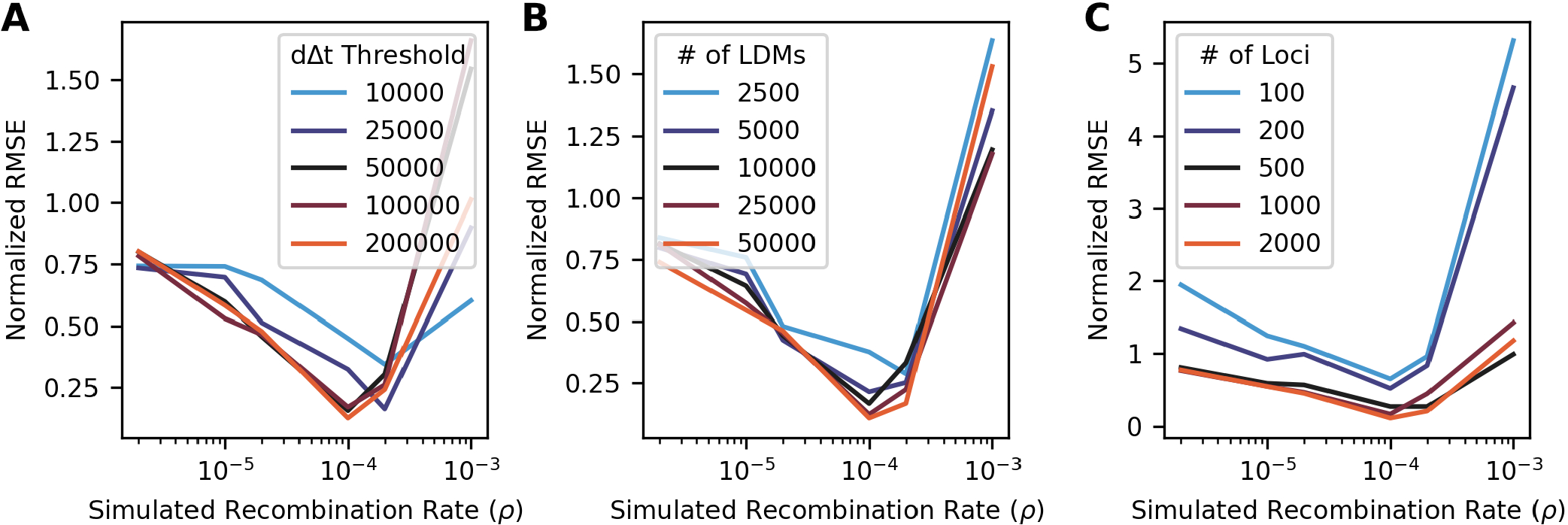
RATS-LD accuracy under varying data set parameters. **(A)** Normalized root mean squared error (RMSE) for RATS-LD estimation when data was truncated at a specified *d*Δ*t* threshold before curve fitting and estimation. Ten simulated data sets were used for the analysis at each *ρ* value with each data set in this panel containing *≈*10, 000 LDMs. **(B)** Normalized RMSEs for 10 simulated data sets with varying numbers of linkage decay measures (LDMs) available for estimation. We downsampled each group by including a LDM in the downsampled data set only if both of its corresponding loci had been randomly drawn. Starting with a dataset containing 0 segregating loci, we drew new segregating loci until the total number of LDMs reached the desired data set size. Error decreases as more LDMs are used for estimation. **(C)** Normalized RMSE for 10 data sets with varying numbers of segregating loci. Segregating loci were sampled so each simulation in the data set contributed an equal number of loci. Estimation error decreases as more segregating loci are used for estimation.

**Figure S12:**
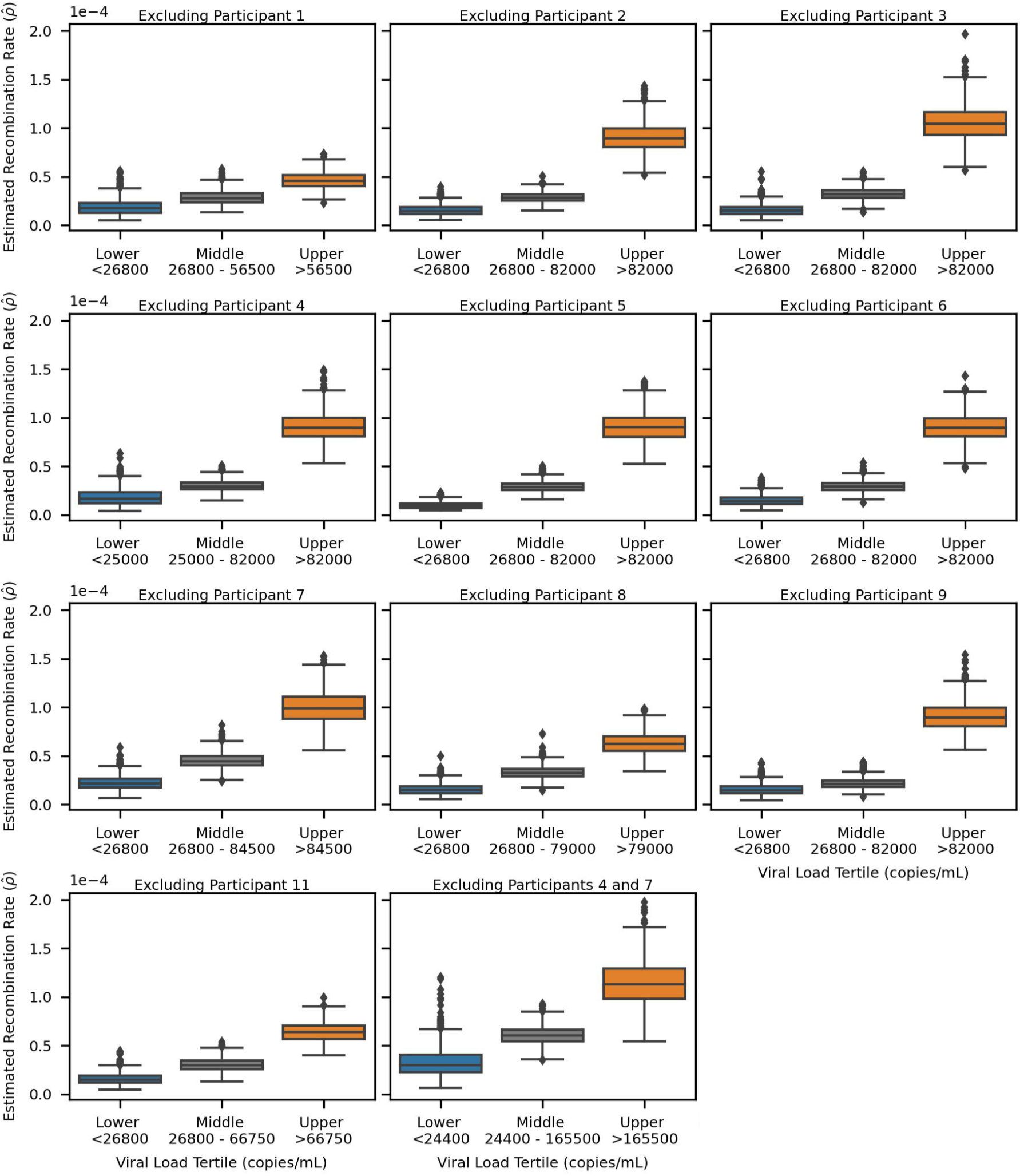
Recombination rate estimates from *in vivo* leave one out analysis also show a positive association between viral load and recombination rate. Each subplot indicates the bootstrapped distributions of recombination rate estimates (1000 Boostraps) when RATS-LD is run on all of the *in vivo* data except for loci pairs from the given participant. An additional panel is included where the data from both participants 4 and 7 are excluded since these individuals were superinfected [56] and show longer range linkage preservation than the other individuals (Figure S6). Each data set was split by three even tertiles to create lower, middle, and upper groups based on viral load. Each box indicates the interquartile range of the distribution.

**Figure S13:**
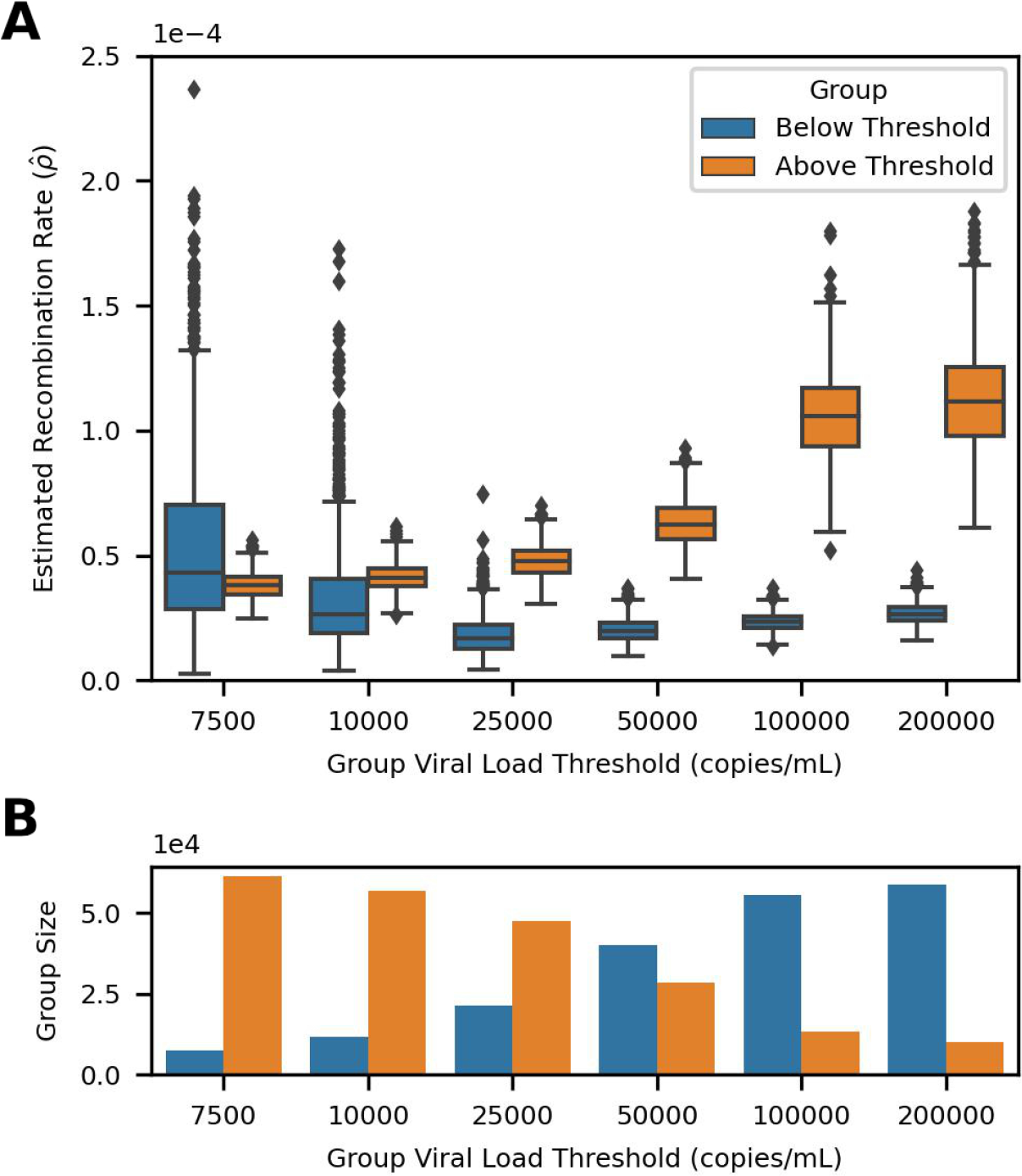
Using a moving threshold to separate viral load groups reveals a dose dependent relationship between viral load and estimated recombination rate 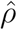. **(A)** Recombination rate estimates, 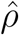, are plotted as a bootstrapped distribution (1000 bootstraps) for groups with varying viral load thresholds separating them. Each box represents the quartiles of the 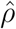 distribution and the median is indicated by the central line. Estimates differ between groups at increasing viral load thresholds. **(B)** The number of LDMs available for estimation in each group is indicated by the height of its corresponding bar.

**Figure S14:**
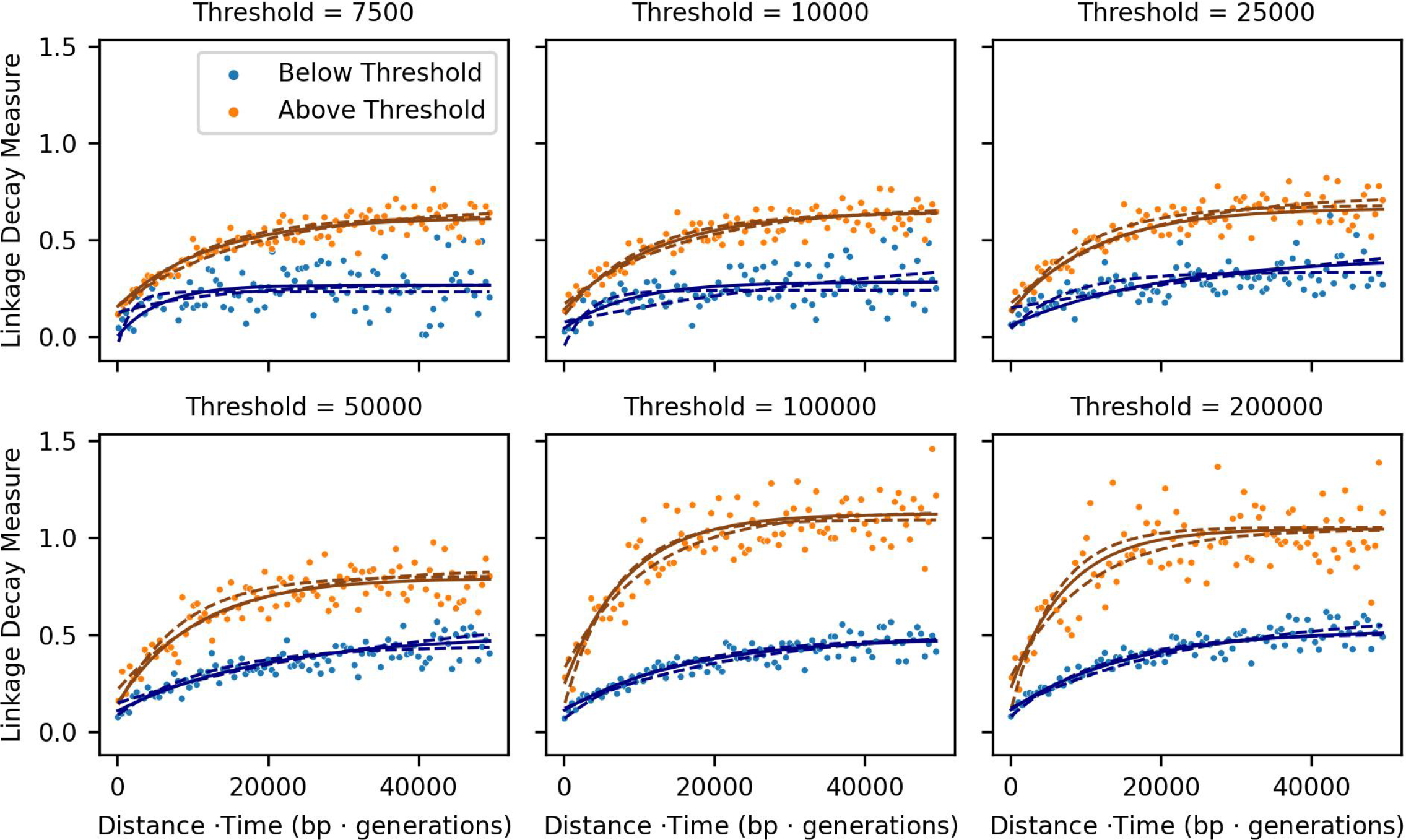
Trends in mean linkage decay measure (LDM) show that linkage decays more rapidly as the viral load threshold (copies/mL) separating low and high viral load groups increases. Each panel contains the linkage decay curves, with binned averages of the LDMs represented by dots, alongside several fits to the data (moving average for display purposes only, window size = 500 bp · generations). The fit producing the median estimate for the bootstrapped distribution (1000 bootstraps) is indicated by a solid line while the fits that produced the estimates at the 0.025 and 0.975 quantiles of the bootstrapped distribution are represented by dashed lines. Panels are separated by the viral load threshold dividing the two groups.

**Figure S15:**
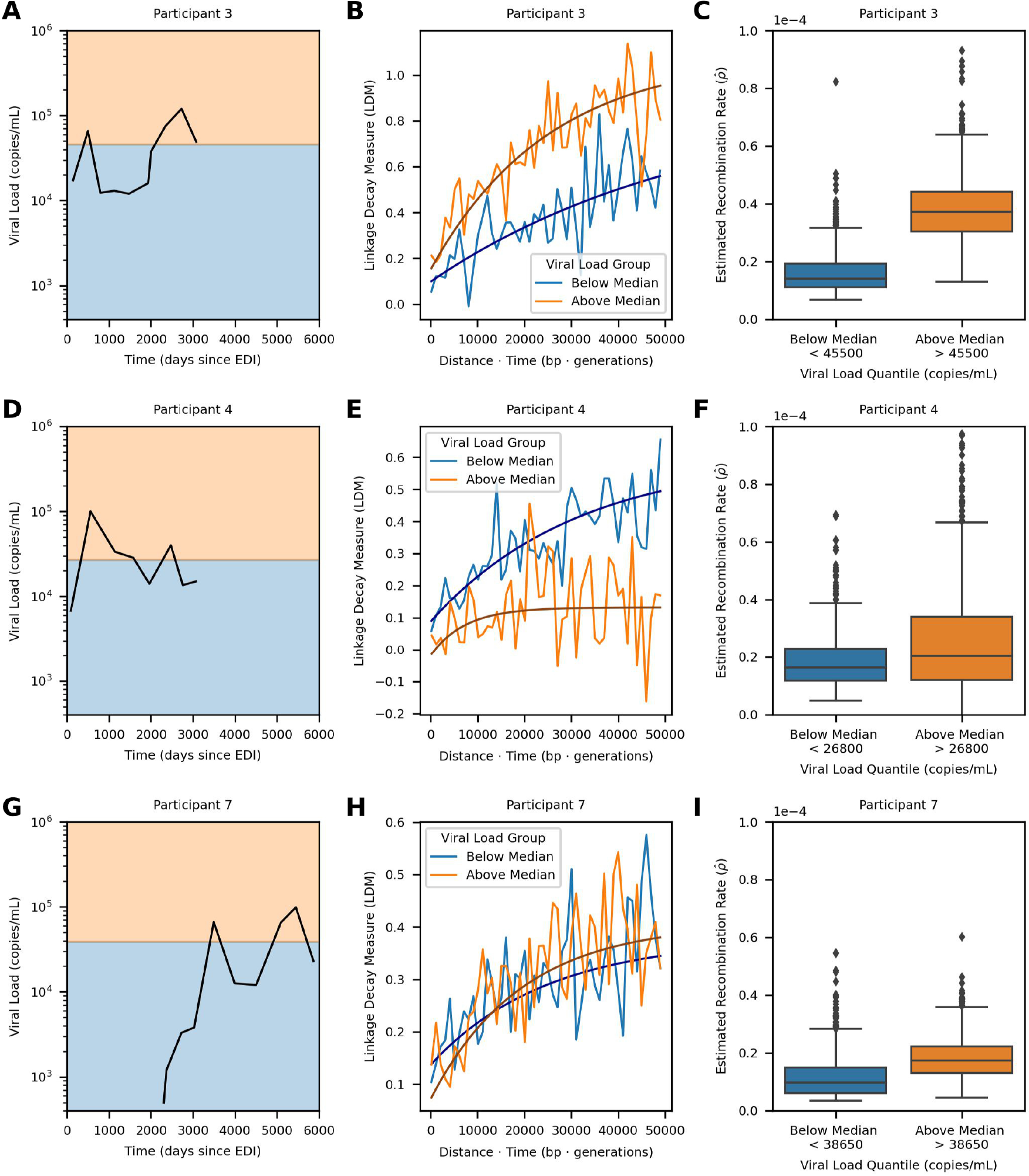
Recombination rate estimation within individuals p3, p4, and p7. Each row shows the results of a RATS-LD analysis run on only data from Participant 3 (top), 4 (middle), or 7 (bottom). The first plot in the row **(A, D, G)** shows the viral load trajectory for the given individual, with background shading indicating the two groups (above median in orange and below median in blue) that pairs of time points were divided into based on their average viral load. **(B, E, H)** RATS-LD curve fits representing the median bootstrapped estimate for each group are plotted on top of the binned average of the LDMs from the given participant’s data (moving average used for display purposes only, window size = 1000 bp ·generations). **(C, F, I)** Box and whisker plots, with each box representing the quartiles of the 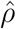 distribution, show the bootstrapped distributions (1000 bootstraps) of recombination rate estimates, 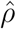, for each individual.

**Figure S16:**
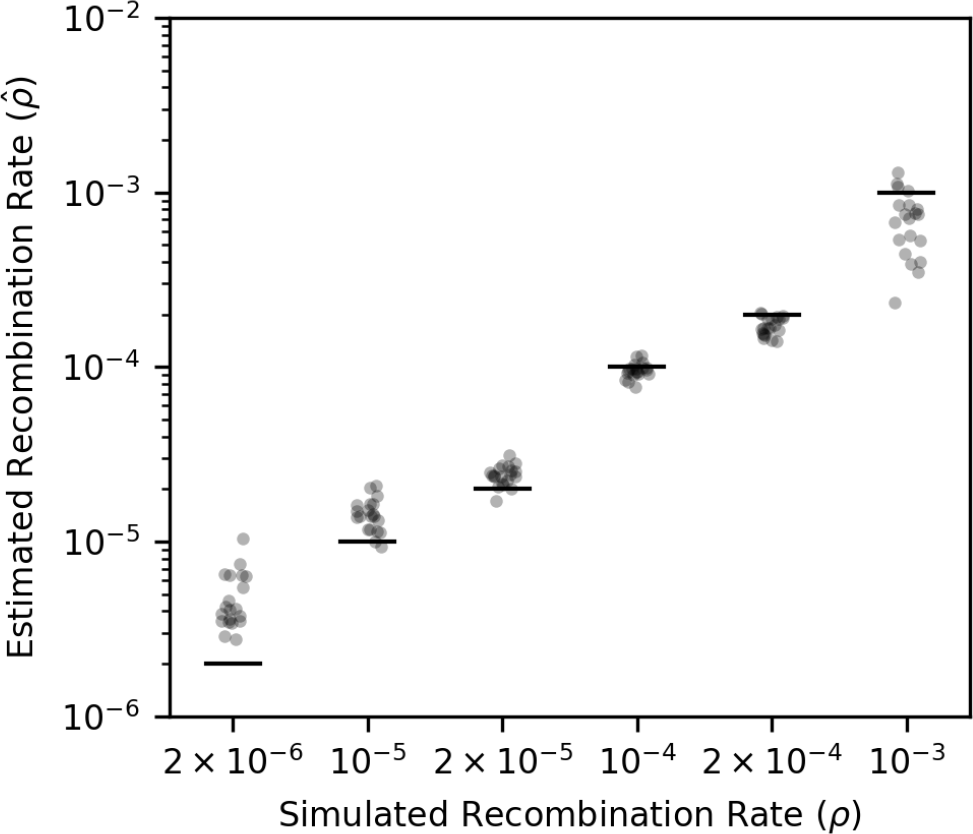
Recombination rate can also be estimated by using the D statistic to measure linkage in simulated data. Each dot represents the median of a bootstrapped estimate distribution (1000 bootstraps). Recombination rate estimates are from RATS-LD runs where the *D* statistic (*D* = *p*_12_ *− p*_1_*p*_2_ where *p*_1_ and *p*_2_ are the frequencies of the the majority alleles at the corresponding loci and *p*_12_ is the haplotype frequency composed of the majority alleles) was used to measure linkage decay rather than the normalized version, *D*′.

**Figure S17:**
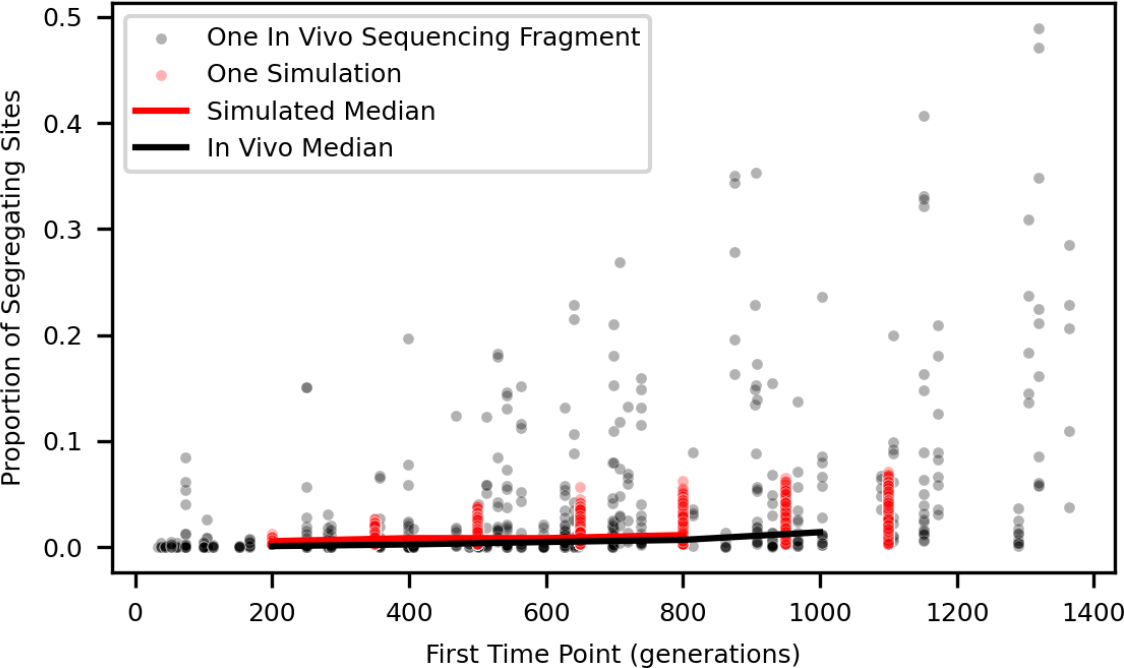
Selection simulations match the proportion of segregating sites observed *in vivo*. The number of segregating sites in simulated and *in vivo* [56] data sets is shown here after data filtering and LDM calculations (See Materials & Methods for details). For each time point pair in these data sets, the proportion of segregating sites (# of sites/sequence length) was plotted against the first time point in the pair. Each dot represents the result for one time point pair in either a simulation (red) or an *in vivo* sequencing fragment (black). Medians were calculated and displayed using 5 bins (bin width = 200 generations).

**Figure S18:**
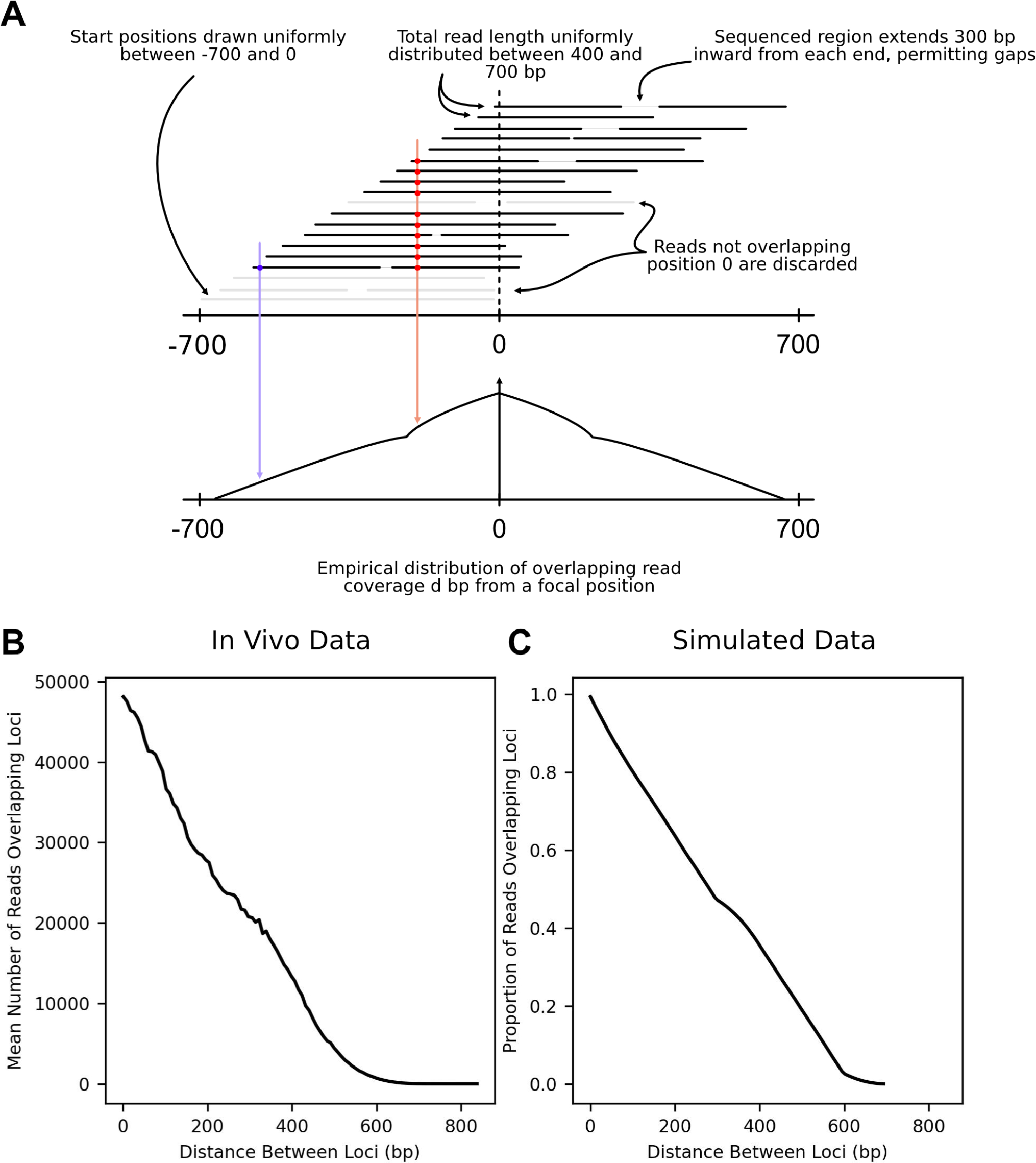
An empirical distribution of read coverage is used to mimic the *in vivo* read coverage simulated data sets. **(A)**. Schematic of how the empirical distribution of read coverage is formed. **(B)** Moving average of the number of reads observed to be overlapping pairs of loci *d* distance apart in the *in vivo* data set (moving average window size = 8bp). **(C)** Moving average of the proportion of reads overlapping loci in the empirical distribution formed via the sampling scheme described in **(A)** (moving average window size = 8bp).

